# Re-evaluation of lysyl hydroxylation in the collagen triple helix: lysyl hydroxylase 1 and prolyl 3-hydroxylase 3 have site-differential and collagen type-dependent roles in lysine hydroxylation

**DOI:** 10.1101/2019.12.16.877852

**Authors:** Yoshihiro Ishikawa, Yuki Taga, Keith Zientek, Nobuyo Mizuno, Antti M. Salo, Olesya Semenova, Sara Tufa, Douglas R. Keene, Paul Holden, Kazunori Mizuno, Johanna Myllyharju, Hans Peter Bächinger

**Affiliations:** Department of Biochemistry and Molecular Biology, Oregon Health & Science University, Portland, Oregon, USA; Shriners Hospital for Children, Research Department, Portland, Oregon, USA; Nippi Research Institute of Biomatrix, Ibaraki, Japan; Oulu Center for Cell-Matrix Research, Biocenter Oulu and Faculty of Biochemistry and Molecular Medicine, University of Oulu, Oulu, Finland

## Abstract

Collagen is the most abundant protein in humans and is heavily post-translationally modified. Its biosynthesis is very complex and requires three different types of hydroxylation (two for proline and one for lysine) that are generated in the rough endoplasmic reticulum (rER). These processes involve many enzymes and chaperones which were collectively termed the molecular ensemble for collagen biosynthesis. However, the function of some of the proteins in this molecular ensemble is controversial. While prolyl 3-hydroxylase 1 and 2 (P3H1, P3H2) are bona fide collagen prolyl 3-hydroxylases, the function of prolyl 3-hydroxylase 3 (P3H3) is less clear. A recent study of P3H3 null mice demonstrated that this enzyme had no activity as prolyl 3-hydroxylase but may instead act as a chaperone for lysyl hydroxylase 1 (LH1). LH1 is required to generate hydroxylysine for crosslinking within collagen triple helical sequences. If P3H3 is a LH1 chaperone that is critical for LH1 activity, P3H3 and LH1 null mice should have similar deficiency in lysyl hydroxylation. To test this hypothesis, we compared lysyl hydroxylation in type I and V collagen from P3H3 and LH1 null mice. Our results indicate LH1 plays a global role for lysyl hydroxylation in triple helical domain of type I collagen while P3H3 is indeed involved in lysyl hydroxylation particularly at crosslink formation sites but is not required for all lysyl hydroxylation sites in type I collagen triple helix. Furthermore, although type V collagen from LH1 null mice surprisingly contained as much hydroxylysine as type V collagen from wild type, the amount of hydroxylysine in type V collagen was clearly suppressed in P3H3 null mice. In summary, our study suggests that P3H3 and LH1 likely have two distinct mechanisms to distinguish crosslink formation sites from other sites in type I collagen and to recognize different collagen types in the rER.

**Author summary:** Collagen is one of the most heavily post-translationally modified proteins in the human body and its post-translational modifications provide biological functions to collagen molecules. In collagen post-translational modifications, crosslink formation on a collagen triple helix adds important biomechanical properties to the collagen fibrils and is mediated by hydroxylation of very specific lysine residues. LH1 and P3H3 show the similar role in lysine hydroxylation for specific residues at crosslink formation sites of type I collagen. Conversely, they have very distinct rules in lysine hydroxylation at other residues in type I collagen triple helix. Furthermore, they demonstrate preferential recognition and modification of different collagen types. Our findings provide a better understanding of the individual functions of LH1 and P3H3 in the rER and also offer new directions for the mechanism of lysyl hydroxylation followed by crosslink formation in different tissues and collagens.

## Introduction

Collagen is not only the most abundant protein, but is also one of the most heavily post-translationally modified proteins in the human body [1, 2]. These post-translational modifications (PTMs) play essential roles in providing biological functions to collagen molecules. Two distinct classifications of PTM exist prior to the incorporation of collagen into extracellular matrices (ECMs), occurring in the unfolded state (a single α-chain) in the rough endoplasmic reticulum (rER) and the folded state (triple helical structure) in the Golgi and the ECM space [3, 4]. Interestingly, there is the case that the extent of the PTM in the Golgi and the ECM space is governed by the PTMs in the rER. Crosslink formation is an important PTM occurring on a collagen triple helical structure and adds important biomechanical properties to the collagen fibrils [5, 6]. However, the pathway of crosslink formation in type I collagen depends on the presence and absence of lysyl hydroxylation in both the collagenous and the telopeptide region [7, 8]. Additionally, *O*-glycosylation, which is generated after lysyl hydroxylation, is involved in crosslink formations and the amount depends on the type of tissue, the rate of triple helix formation and the presence or absence of ER chaperones [9–12]. Thus, PTMs of unfolded α-chains in the rER are critical for quality control in a collagen ultrastructure.

Collagen biosynthesis including PTMs is complex and involves many enzymes and chaperones, which are collectively termed the molecular ensemble [13]. Because some of the enzymes only can modify unfolded, and not triple helical, collagen chains the time that collagen chains remain unfolded in the rER is a critical factor for correct PTMs and foldases control the rate of triple helix formation [14–17]. There are three hydroxylations (proline 3-hydroxylation, proline 4-hydroxylation and lysine hydroxylation) that are fundamental PTMs occurring before triple helix formation [3, 13]. Interestingly, the function of prolyl 3-hydroxylase 3 (P3H3) is controversial. It has been suggested that this protein has no prolyl hydroxylase activity and instead acts as a chaperone for lysyl hydroxylase 1 (LH1) [18]. LHs hydroxylate specific lysine residues in both the collagenous and telopeptide regions, and the three isoforms (LH1, 2 and 3) have been proposed to play specific roles based on collagen sequences [16]. LH2 is specific for hydroxylating the telopeptide [16], and LH1 has been suggested to hydroxylate the triple-helical regions [13, 19–21]. The study of patients with LH1 mutations, and of LH3 mutant mice indicate that both LH1 and LH3 could have substrate preferences (e.g. LH1 and LH3 prefer type I/III collagen and type II/IV/V collagen, respectively) [22–25]. However, this indication could not fully explain the diversity of lysyl hydroxylation in tissues of the LH1 null mouse model and type I collagen from different tissues of Ehlers-Danlos Syndrome (EDS)-VIA patients [23, 26]. Few analyses have investigated the level of lysyl hydroxylation in purified collagens from mutant or LH knockout models using both qualitative and quantitative measurements.

In this study, we aimed to re-evaluate the role of LH1 in collagen triple helices and test whether P3H3 is essential for the LH1 activity. To achieve this, we compared the levels of overall lysyl hydroxylation and PTM occupancy at individual sites between collagens extracted from P3H3 or LH1 null mice. If P3H3 is a chaperone required for LH1 activity as recently suggested [18], then both P3H3 and LH1 null mice should cause similar defects in lysine hydroxylation. Moreover we analyzed different collagens from different tissues to test the hypothesis that LH1 is a collagen type-specific enzyme [22–25], and to understand if P3H3 might contribute to this differential activity. This direct comparison of different collagens from different tissues from these two null mouse lines provide a better understanding of the individual functions of P3H3 and LH1 in the rER.

## Results

### Basic characterization of P3H3 null mice

P3H3 null mice were generated by Ozgene as shown in Figure 1A and with more detailed information in the methods section. Figure 1B displays the result of PCR genotyping showing that the P3H3 null allele product is smaller than WT due to the deletion of Exon 1. To confirm that the expression of P3H3 protein was abolished, Western blotting was performed using a whole kidney lysate and the protein signal corresponding to MW of P3H3 (79 kDa) was absent in the P3H3 null lysate (Figure 1C). As previously reported [18], P3H3 null mice were also viable and we did not observe any obvious growth or skeletal phenotypes by growth curves and X-ray images, respectively (Figure 1D and E). LH1 null mice were generated and characterized previously [26].

**Figure 1.**
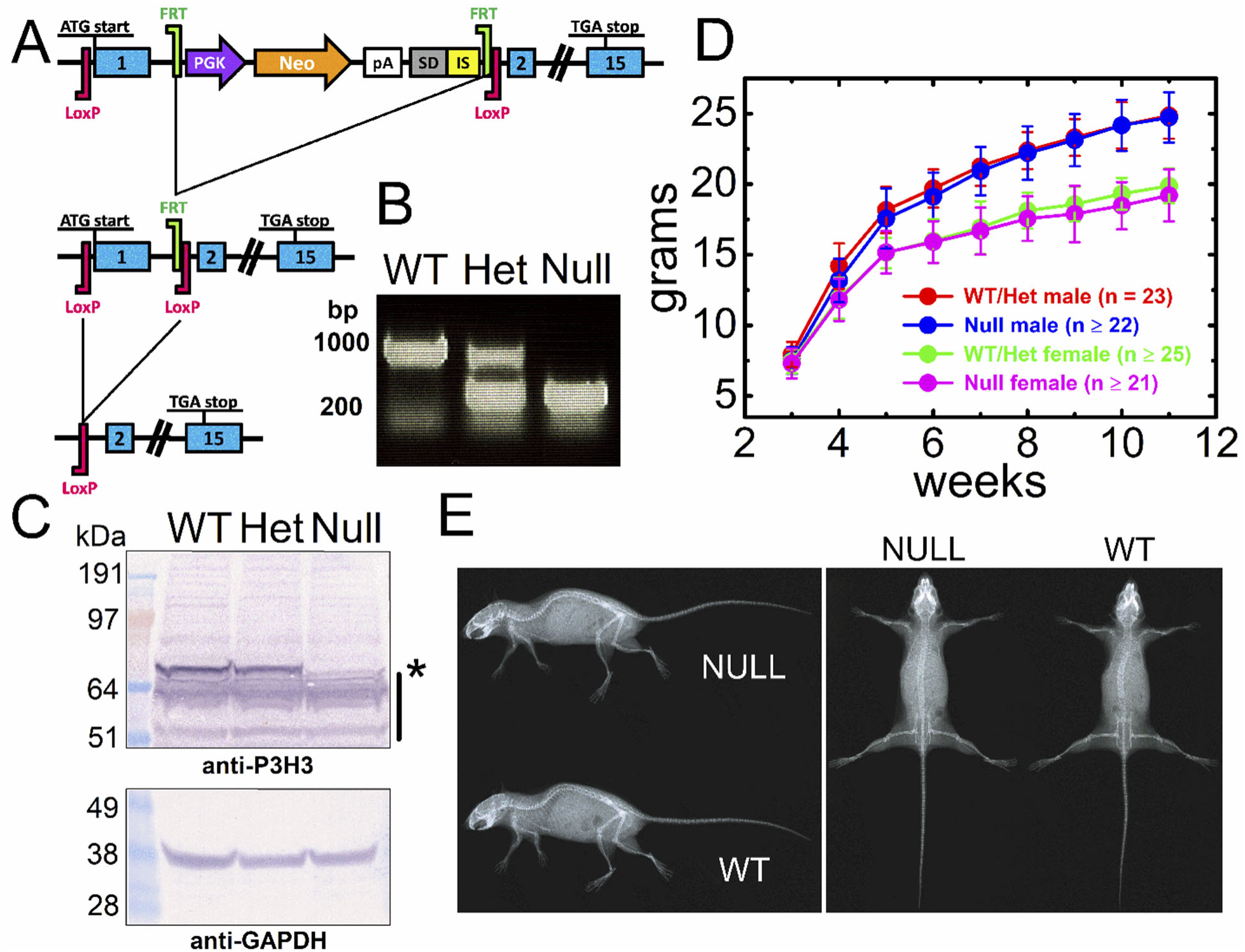
Generation of P3H3 null mice. (A) Strategy for generation of the P3H3 null mice. The mouse *Leprel2* gene was eliminated using FRT sites and loxP sites by an FLPe recombinase followed by a Cre recombinase in ES cell, respectively. (B) PCR genotyping of P3H3 wild type (WT), heterozygote (Het) and homozygote (Null) mice and smaller PCR products were generated in Het and Null due to deletion of Exon 1. (C) Total soluble proteins from 3-month-old whole mouse kidney in P3H3 WT, Het and Null were extracted and blotted by using anti-GAPDH and anti-P3H3. The asterisk and black line in the panel of P3H3 indicate the band of P3H3 and non-specific bands generated by antibodies, respectively. (D) Whole body weights of P3H3 WT and Het male (red) and female (green) and P3H3 Null male (blue) and female (magenta) were plotted as a function of age with standard deviations. Growth differences were not observed. (E) Full body x-rays of 4-month-old female P3H3 WT and Null were scanned from top and side angles. No skeletal defects were found.

### Biochemical characterization of purified type I collagen from different tissues of P3H3 and LH1 null mouse models

To enable qualitative and quantitative analyses, type I collagen was purified from tissues by pepsin treatment followed by sodium chloride precipitation. We analyzed three different tissues (tendon, skin and bone) from each mouse model. We evaluated the level of PTMs by comparing migration using SDA-PAGE [27] and determined the thermal stability using circular dichroism (CD) spectra [28, 29] of purified type I collagen from the different tissues of P3H3 null and LH1 null mice (Figure 2). Type I collagen from both P3H3 null and LH1 null skin migrates a little faster and shows a lower melting temperature than WT, whereas there is no clear difference for type I collagen from tendon and bone between WT and nulls in gel migration or melting temperature (Figure 2). This suggests that skin is the most affected tissue in both P3H3 null and LH1 null mice.

**Figure 2.**
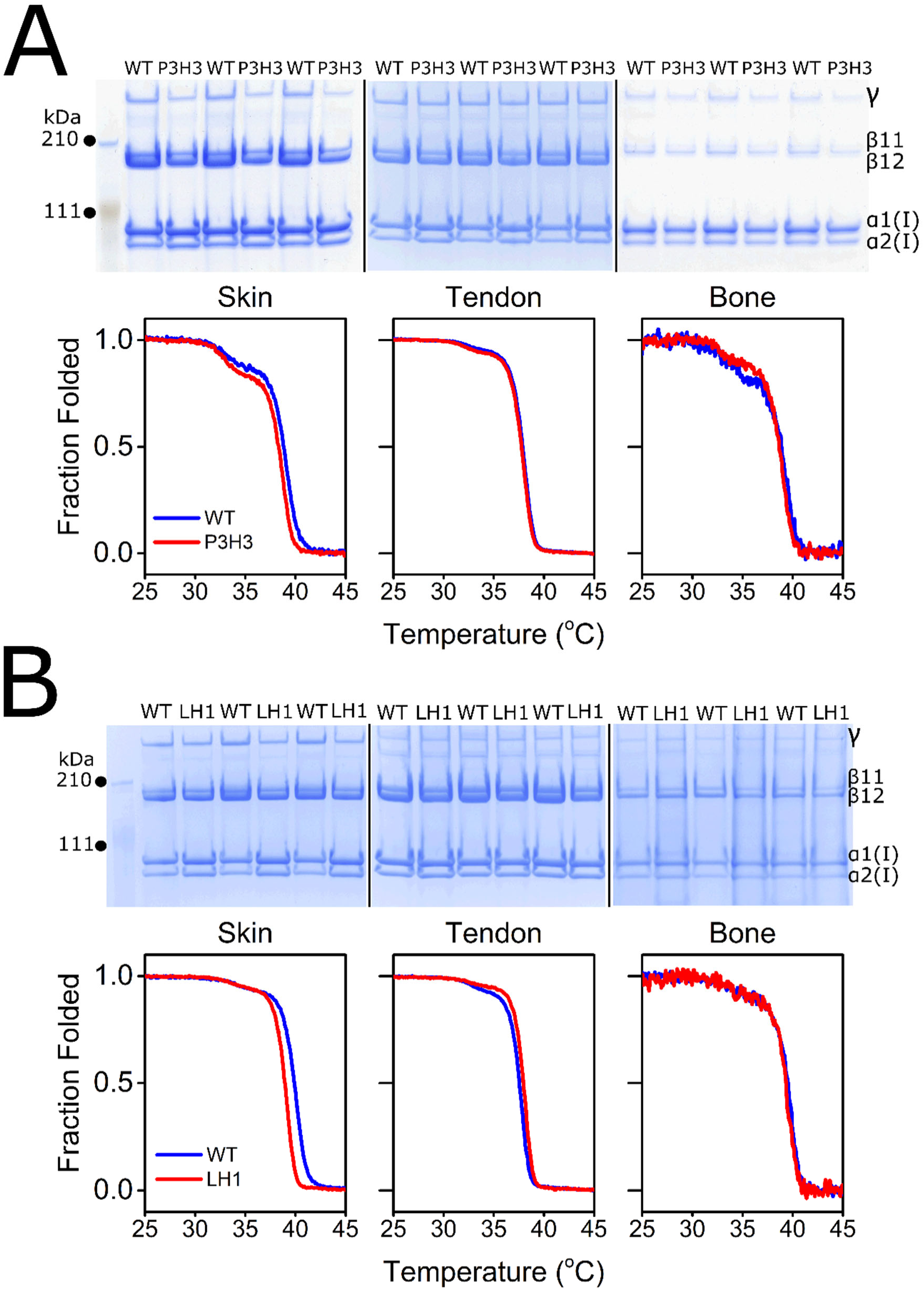
Biochemical characterization of type I collagen from P3H3 and LH1 Null mice. (A: upper panel) SDS-PAGE analysis of purified pepsin treated type I collagen of P3H3 WT and null from skin, tendon and bone. Figure shows the final purified material in the presence of reducing agent running on a NuPAGE 3 – 8 % Tris-Acetate gel (ThermoFisher) stained with GelCode Blue Stain Reagent (ThermoFisher). (A: lower panel) Thermal stability of type I collagen of P3H3 WT (blue) and null (red) from skin, tendon and bone were monitored by CD at 221 nm in 0.05 M acetic acid at 10 °C/h heating rate. P3H3 indicates P3H3 null and tissues were collected from 2∼5-month-old mice. Biological replicates of each curve were n = 3 for skin P3H3 null, tendon P3H3 WT and null, n = 4 for skin P3H3 WT and bone P3H3 WT and n = 5 for bone P3H3 null. (B: upper panel) SDS-PAGE analysis of purified pepsin treated type I collagen of LH1 WT and null from skin, tendon and bone. Figure shows the final purified material in the presence of reducing agent running on a NuPAGE 3 – 8 % Tris-Acetate gel (ThermoFisher) stained with GelCode Blue Stain Reagent. (B: lower panel) Thermal stability of type I collagen of LH1 WT (blue) and null (red) from skin, tendon and bone were monitored by CD at 221 nm in 0.05 M acetic acid at 10 °C/h heating rate. LH1 indicates LH1 null and tissues were collected from 10-week-old mice. Biological replicates of each curve were n = 4 for all tissues and genotypes. For both SDS-PAGE analysis of P3H3 and LH1, each genotype had three biological replicates since each lane in gel was loaded by independently prepared collagen from tissue.

### Quantitative analysis to determine the total level of post-translational modifications of type I collagen in P3H3 null and LH1 null mice

Amino acid analysis (AAA) was used to quantify the total number of PTMs in the purified type I collagens. Neither P3H3 nor LH1 null mice had changes in proline hydroxylations (prolyl 3- and 4-hydroxylation), however, both strains had interesting changes in lysyl hydroxylation (Figure 3 and Table 1). LH1 deficiency significantly decreased the amounts of hydroxylysine in tendon, skin and bone although bone was a slightly lesser extent. In contrast, P3H3 deficiency had a much smaller effect on lysyl hydroxylation than LH1 whereby skin showed further reduction of hydroxylysine compared to tendon and bone. Next, we determined the occupancy of *O*-glycosylation of hydroxylysine in tendon and skin by liquid chromatography–mass spectrometry (LC– MS). The calculated value of galactosyl hydroxylysine (GHL) does not show any significant difference in skin but does slightly increase in tendon for both P3H3 null and LH1 null mice (Figure 4 and Table 2). Interestingly, the magnitude of change in unmodified hydroxylysine and glucosylgalactosyl hydroxylysine (GGHL) seems to have some correlation in both tendon and skin. For example, both P3H3 null and LH1 null type I collagen in tendon showed unmodified hydroxylysine was decreased whereas GGHL was increased by a similar magnitude of decreasing unmodified hydroxylysine (Figure 4 and Table 2). While the effect is opposite manner, the same observation is also seen in skin of P3H3 null and LH1 null type I collagen (Figure 4 and Table 2). This indicates that the affected site(s) in the absence of P3H3 and LH1 are different location or have different level of *O*-glycosylation between tendon and skin. Taken together, P3H3 clearly has a role in lysyl hydroxylation, but the level of lysyl hydroxylation in type I collagen is more drastically decreased in LH1 null mice compared to P3H3 null mice. Furthermore, the data also suggests that P3H3 and LH1 likely have distinct roles in lysyl hydroxylation and sugar attachment between tissues.

**Figure 3.**
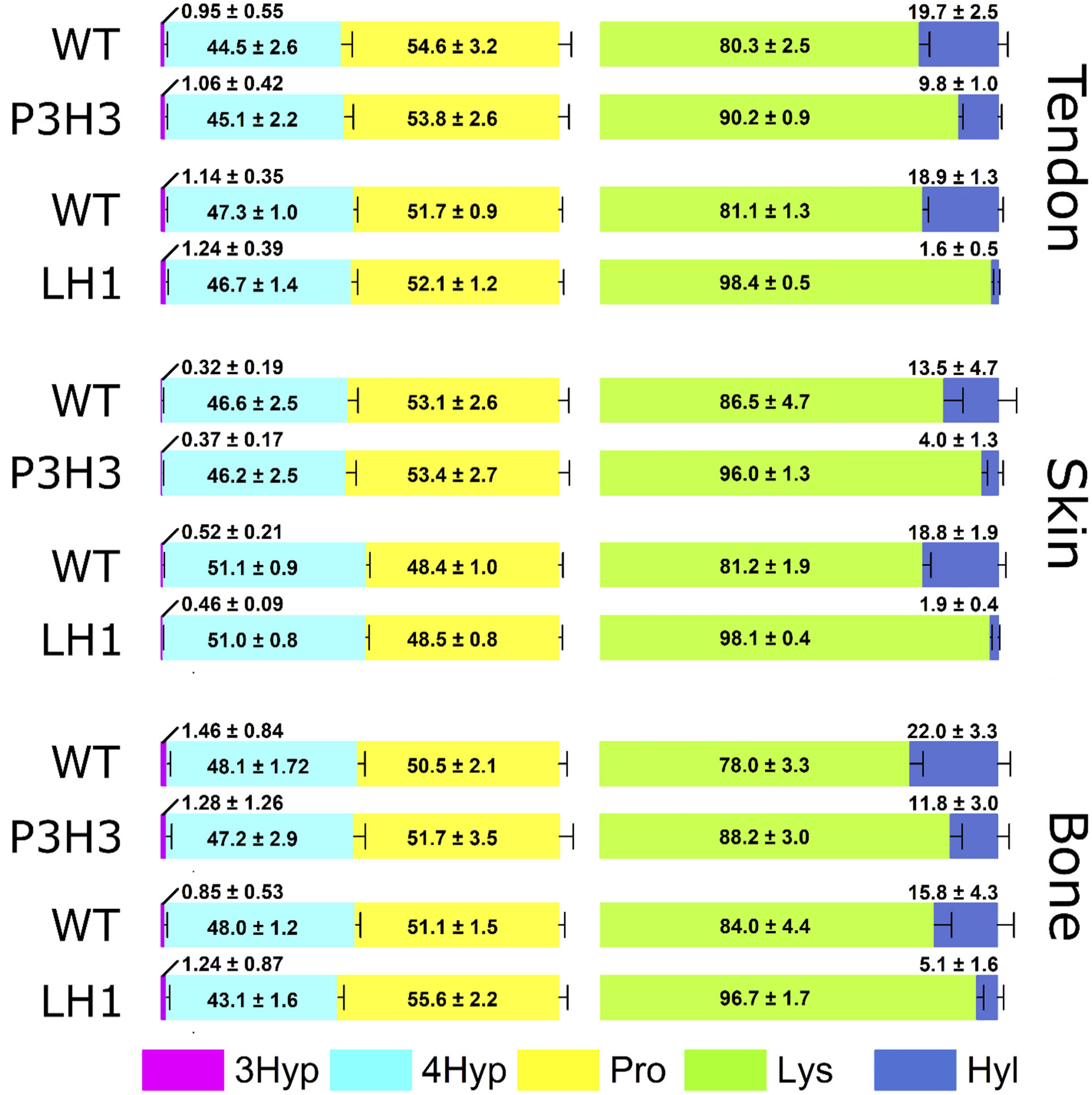
Summary of overall proline and lysine post-translational modifications in type I collagen between tissues. The ratio of post-translational modifications in proline (3Hyp + 4Hyp + Pro = 100) and lysine (Lys + Hyl = 100) in type I collagen of WTs, P3H3 null and LH1 null from tendon, skin and bone are demonstrated as bar graphs which are generated by values from Table 1. Values of amino acids were obtained using amino acid analysis and biological replicates were tendon: n ≥ 8, skin: n = 8 and bone n ≥ 4. The numbers in the graphs indicate the mean ± S.D. of individual amino acids and P values obtained by statistical analyses are in Table 1 [3Hyp(magenta); 3-hydroxyproline, 4Hyp (cyan); 4-hydroxyproline, Pro (yellow); unmodified proline, Lys (green); unmodified lysine, Hyl (blue); hydroxylysine]. P3H3 and LH1 indicate P3H3 null and LH1 null, respectively.

**Figure 4.**
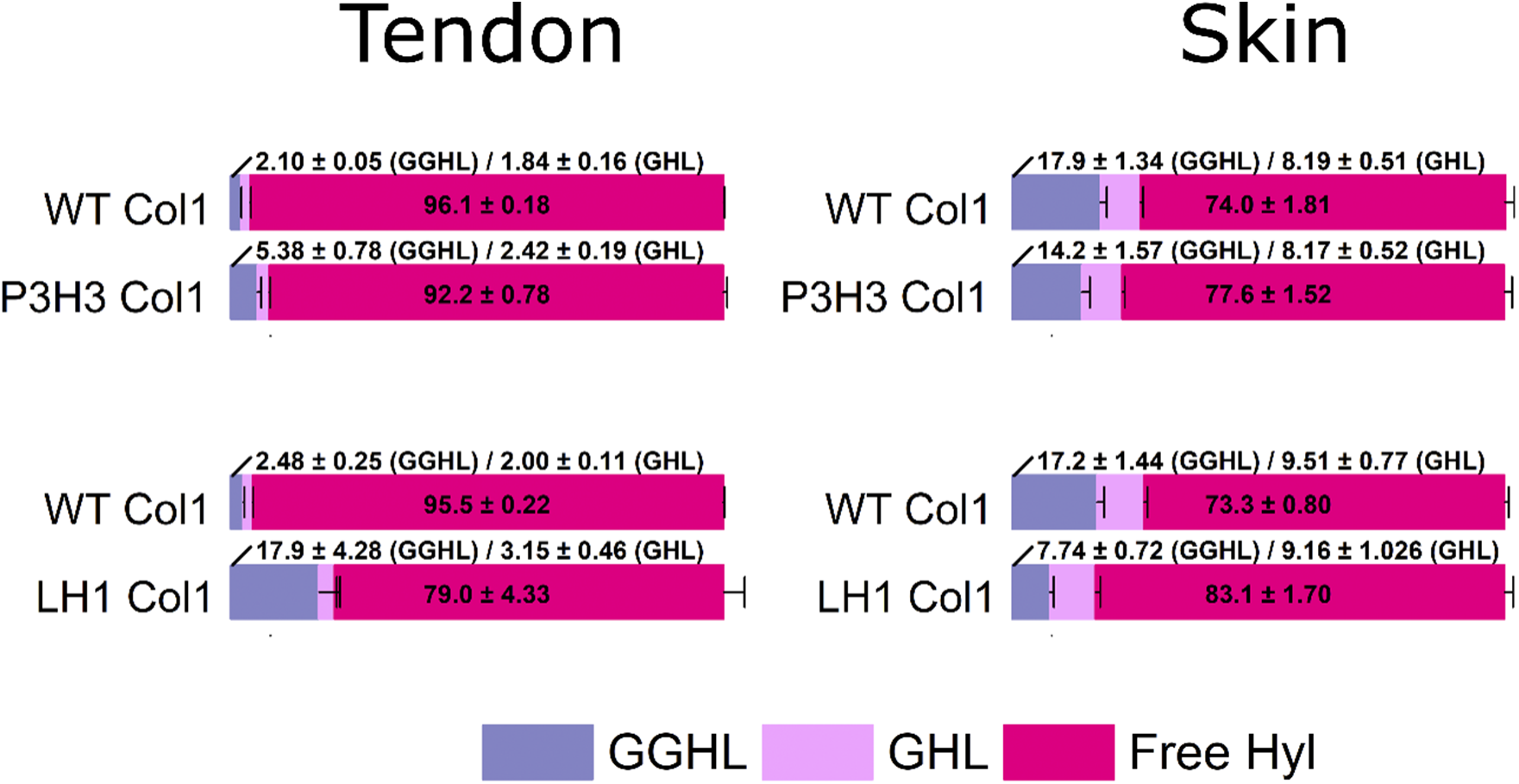
Summary of overall occupancy of *O*-glycosylation attached to hydroxylysine in type I collagen between tissues. The occupancy of *O*-glycosylation attached to hydroxylysine (GGHL + GHL + Free Hyl = 100) in type I collagen of WTs, P3H3 null and LH1 null from tendon and skin are demonstrated as bar graphs which are generated by values from Table 2. Values of *O*-glycosylation and unmodified hydroxylysine were obtained by LC–MS after alkaline hydrolysis and biological replicates were n = 4 for all tissues and genotypes. The numbers in the graphs indicate the mean ± S.D. of individual *O*-glycosylation and unmodified hydroxylysine and P values obtained by statistical analyses are in Table 2 [GGHL (gray); glucosylgalactosyl hydroxylysine, GHL (light pink); galactosyl hydroxylysine, Free Hyl (dark pink); unmodified hydroxylysine]. In figure, P3H3 and LH1 indicate P3H3 null and LH1 null, respectively.

**Table 1:** Comparison of overall proline and lysine post-translational modifications in type I collagen between tissues and type V collagen in skin.

|  |  |  | 3Hyp (%) | P value | 4Hyp (%) | P value | Pro (%) | P value | Lys (%) | P value | Hyl (%) | P value |
| --- | --- | --- | --- | --- | --- | --- | --- | --- | --- | --- | --- | --- |
| col1 | tendon | WT | 0.95 ± 0.55 | 0.65658 | 44.5 ± 2.6 | 0.60898 | 54.6 ± 3.2 | 0.61666 | 80.3 ± 2.5 | 5.36786E-8 | 19.7 ± 2.5 | 5.36786E-8 |
|  |  | P3H3 | 1.06 ± 0.42 |  | 45.1 ± 2.2 |  | 53.8 ± 2.6 |  | 90.2 ± 0.9 |  | 9.8 ± 1.0 |  |
|  |  | WT | 1.14 ± 0.35 | 0.5239 | 47.3 ± 1.0 | 0.21519 | 51.7 ± 0.9 | 0.25603 | 81.1 ± 1.3 | 0 | 18.9 ± 1.3 | 0 |
|  |  | LH1 | 1.24 ± 0.39 |  | 46.7 ± 1.4 |  | 52.1 ± 1.2 |  | 98.4 ± 0.5 |  | 1.6 ± 0.5 |  |
|  | skin | WT | 0.32 ± 0.19 | 0.61795 | 46.6 ± 2.5 | 0.70334 | 53.1 ± 2.6 | 0.75642 | 86.5 ± 4.7 | 7.7608E-5 | 13.5 ± 4.7 | 7.7608E-5 |
|  |  | P3H3 | 0.37 ± 0.17 |  | 46.2 ± 2.5 |  | 53.4 ± 2.7 |  | 96.0 ± 1.3 |  | 4.0 ± 1.3 |  |
|  |  | WT | 0.52 ± 0.21 | 0.41975 | 51.1 ± 0.9 | 0.80229 | 48.4 ± 1.0 | 0.71858 | 81.2 ± 1.9 | 8.29004E-13 | 18.8 ± 1.9 | 8.29004E-13 |
|  |  | LH1 | 0.46 ± 0.09 |  | 51.0 ± 0.8 |  | 48.5 ± 0.8 |  | 98.1 ± 0.4 |  | 1.9 ± 0.4 |  |
|  | bone | WT | 1.46 ± 0.84 | 0.79823 | 48.1 ± 1.72 | 0.59762 | 50.5 ± 2.1 | 0.50736 | 78.0 ± 3.3 | 8.81801E-4 | 22.0 ± 3.3 | 8.81801E-4 |
|  |  | P3H3 | 1.28 ± 1.26 |  | 47.2 ± 2.9 |  | 51.7 ± 3.5 |  | 88.2 ± 3.0 |  | 11.8 ± 3.0 |  |
|  |  | WT | 0.85 ± 0.53 | 0.43429 | 48.0 ± 1.2 | 0.00101 | 51.1 ± 1.5 | 0.00811 | 84.0 ± 4.4 | 3.94453E-5 | 15.8 ± 4.3 | 3.04315E-5 |
|  |  | LH1 | 1.24 ± 0.87 |  | 43.1 ± 1.6 |  | 55.6 ± 2.2 |  | 96.7 ± 1.7 |  | 5.1 ± 1.6 |  |
| Col5 | skin | WT | 1.12 ± 1.06 | 0.84309 | 44.5 ± 3.0 | 0.99626 | 54.4 ± 3.2 | 0.94792 | 39.6 ± 4.7 | 7.65287E-5 | 60.4 ± 4.7 | 7.65284E-5 |
|  |  | P3H3 | 1.06 ± 0.77 |  | 44.5 ± 3.2 |  | 54.4 ± 2.9 |  | 48.6 ± 6.7 |  | 51.4 ± 6.7 |  |
|  |  | WT | 1.07 ± 0.20 | 0.1898 | 52.5 ± 2.1 | 0.05353 | 46.5 ± 2.3 | 0.03944 | 40.5 ± 2.6 | 0.41188 | 60.0 ± 2.6 | 0.41188 |
|  |  | LH1 | 2.03 ± 1.10 |  | 54.4 ± 0.5 |  | 43.5 ± 1.3 |  | 38.0 ± 4.2 |  | 62.0 ± 4.2 |  |

Values are given as means ± S.D. Biological replicates were tendon: n ≥ 8, bone n ≥ 4, skin type I collagen: n = 8 and skin type V collagen: n ≥ 3.
3Hyp + 4Hyp + Pro = 100 % and Lys + Hyl = 100 %. Values of amino acids were obtained using amino acid analysis.
Note: 3Hyp; 3-hydroxyproline, 4Hyp; 4-hydroxyproline, Pro; unmodified proline, Lys; unmodified lysine, Hyl; hydroxylysine.

**Table 2.**
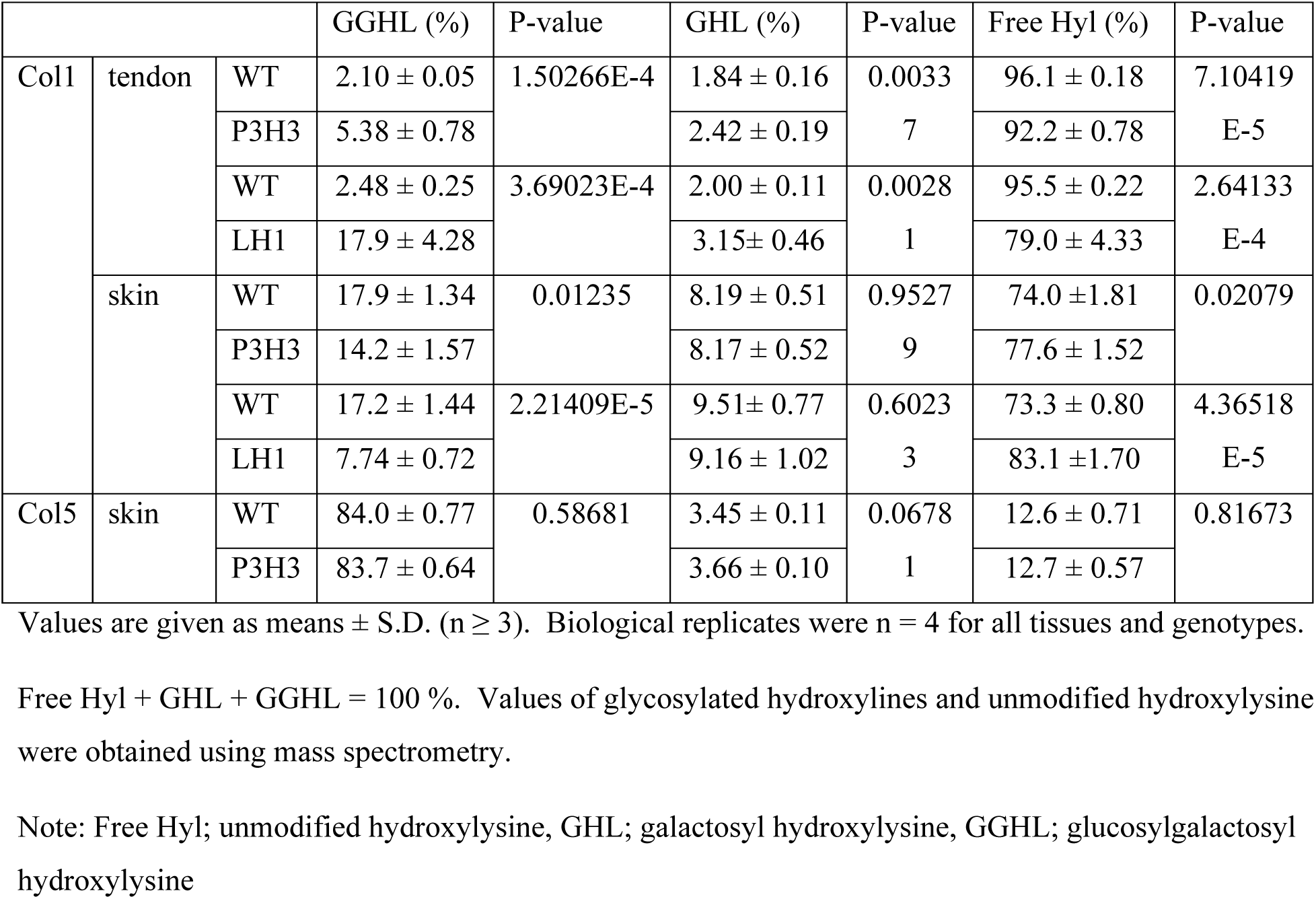
Occupancy (%) of glycosylation in total hydroxylysine residues.

### Individual lysyl hydroxylation site analysis of type I collagen from P3H3 null and LH1 null tissues

The α1 and α2 chain of type I collagen contain multiple lysyl hydroxylation sites in their triple helical sequences [30–32]. A previous report of P3H3 null mice [18] only demonstrated lysyl hydroxylation sites on lysine-87 (K87) in both the α1 and α2 chain. These residues at the amino-terminus of triple helical domain are important for crosslink formation in the ECM [18]. We determined the occupancy of PTMs at individual lysyl hydroxylation sites between WTs, P3H3 and LH1 nulls in tendon and skin (Figure 5, Table 3 and Table 4). In LH1 null tissues, the level of lysyl hydroxylation and subsequent *O*-glycosylation were significantly decreased at all lysyl hydroxylation sites of both tendon and skin. In P3H3 null tissues, we confirmed the reduction of lysine modifications in K87 in both the α1 and α2 of type I collagen as previously reported [18], and a large reduction was also found at α1 K930 and α2 K933 which are near the carboxy-terminus of the triple helical domain involved in crosslink formation. The other sites α1 K99, α1 K174, α2 K174 and α2 K219 also showed clear reduction, however, there was no notable decrease in the level of lysyl hydroxylation at the sites in the middle of the triple helix of α1 chain (α1 K219 and α1 K564). In summary, LH1 might play a global role for lysyl hydroxylation at all sites in the triple helical domain of type I collagen whereas the role of P3H3 could be restricted to specific sites.

**Figure 5.**
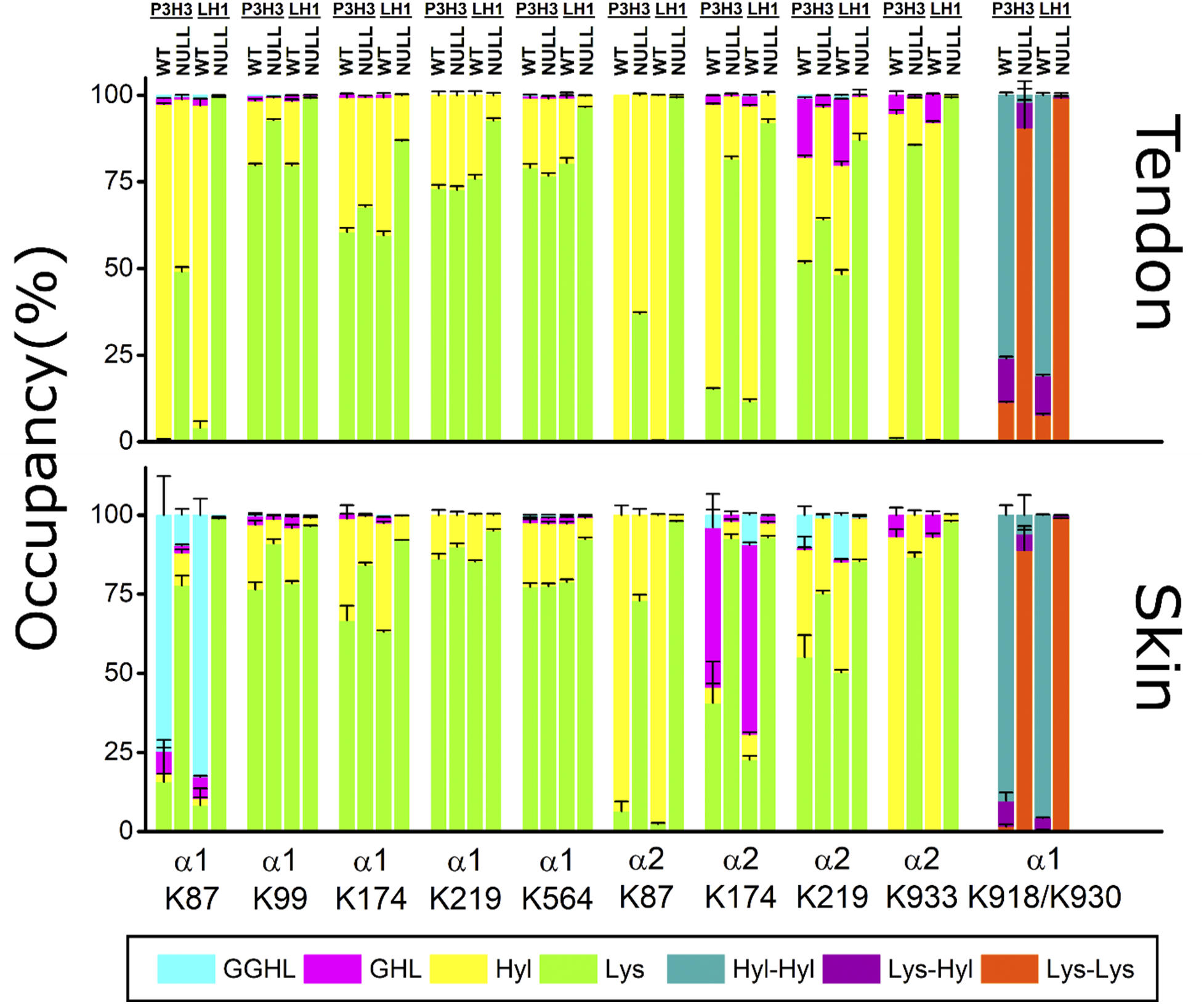
Summary of lysine post-translational modifications of tendon type I collagen in individual sites. Bar graphs represent the occupancy of lysine modifications in individual lysyl hydroxylation sites of type I collagen of WTs, P3H3 null and LH1 from tendon and skin. Individual values [GGHL (cyan); glucosylgalactosyl hydroxylysine, GHL (magenta); galactosyl hydroxylysine, Hyl (yellow); unmodified hydroxylysine, Lys (green); unmodified lysine, Hyl-Hyl (dark cyan); unmodified hydroxylysine and unmodified hydroxylysine, Lys-Hyl (purple); unmodified lysine and unmodified hydroxylysine, Lys-Lys (orange); unmodified lysine and unmodified lysine] correspond to Table 3 and Table 4 for tendon and skin, respectively. Values of modified and unmodified hydroxylines and unmodified lysine were obtained using mass spectrometry and biological replicates were n = 4 for all tissues and genotypes. α1, α2 and K + numbers indicate α1 and α2 chain of type I collagen and residue number from the first residue of triple helical domain.

**Table 3:** Comparison of lysine post-translational modifications of tendon type I collagen at individual sites.

|  |  |  | Lys (%) | P value | Hyl (%) | P value | GHL (%) | P value | GGHL (%) | P value |  |  |  |  |
| --- | --- | --- | --- | --- | --- | --- | --- | --- | --- | --- | --- | --- | --- | --- |
| α1 K87 | P3H3 | WT | 0.56 ± 0.17 | 3.91E-10 | 96.8 ± 0.26 | 4.19E-10 | 1.70 ± 0.07 | 4.84E-07 | 0.93 ± 0.02 | 7.49E-07 | | | | |
|  |  | Null | 49.1 ± 1.29 |  | 49.7 ± 1.25 |  | 0.77 ± 0.04 |  | 0.46 ± 0.04 |  |  |  |  |  |
|  | LH1 | WT | 4.03 ± 1.86 | 5.84E-11 | 93.1 ± 1.85 | 6.77E-11 | 1.76 ± 0.02 | 1.70E-11 | 1.10 ± 0.02 | 1.12E-09 |  |  |  |  |
|  |  | Null | 99.4 ± 0.13 |  | 0.45 ± 0.11 |  | 0.04 ± 0.01 |  | 0.12 ± 0.02 |  |  |  |  |  |
| α2 K87 | P3H3 | WT | 0.35 ± 0.04 | 6.42E-12 | 99.7 ± 0.04 | 6.42E-12 | ND | | ND | | | | | |
|  |  | Null | 36.9 ± 0.49 |  | 63.1 ± 0.49 |  | ND |  | ND |  |  |  |  |  |
|  | LH1 | WT | 0.30 ± 0.07 | 0 | 99.7 ± 0.07 | 0 | ND |  | ND |  |  |  |  |  |
|  |  | Null | 99.3 ± 0.12 |  | 0.66 ± 0.12 |  | ND |  | ND |  |  |  |  |  |
| α1 K99 | P3H3 | WT | 76.8 ± 0.48 | 8.47E-09 | 18.6 ± 0.47 | 1.18E-08 | 1.47 ± 0.02 | 5.96E-10 | 0.10 ± 0.01 | 1.02E-04 | | | | |
|  |  | Null | 92.8 ± 0.33 |  | 6.53 ± 0.33 |  | 0.60 ± 0.01 |  | 0.05 ± 0.01 |  |  |  |  |  |
|  | LH1 | WT | 79.7 ± 0.56 | 7.94E-10 | 18.6 ± 0.53 | 1.05E-09 | 1.59 ± 0.08 | 2.31E-08 | 0.12 ± 0.01 | 1.33E-06 |  |  |  |  |
|  |  | Null | 99.1 ± 0.18 |  | 0.80 ± 0.17 |  | 0.08 ± 0.07 |  | 0 |  |  |  |  |  |
| α1 K174 | P3H3 | WT | 60.5 ± 1.20 | 3.13E-05 | 38.8 ± 1.18 | 3.43E-05 | 0.71 ± 0.03 | 2.22E-04 | ND | | | | | |
|  |  | Null | 67.7 ± 0.51 |  | 31.7 ± 0.54 |  | 0.53 ± 0.03 |  | ND |  |  |  |  |  |
|  | LH1 | WT | 59.5 ± 1.31 | 1.53E-08 | 39.8 ± 1.31 | 1.81E-08 | 0.76 ± 0.05 | 6.70E-08 | ND |  |  |  |  |  |
|  |  | Null | 86.8 ± 0.32 |  | 13.2 ± 0.32 |  | 0 |  | ND |  |  |  |  |  |
| α2 K174 | P3H3 | WT | 15.2 ± 0.26 | 2.95E-12 | 82.2 ± 0.17 | 2.90E-12 | 2.53 ± 0.10 | 8.17E-09 | 0.06 ± 0.01 | 7.01E-05 | | | | |
|  |  | Null | 81.6 ± 0.74 |  | 18.2 ± 0.74 |  | 0.22 ± 0.03 |  | 0 |  |  |  |  |  |
|  | LH1 | WT | 11.6 ± 0.57 | 7.91E-12 | 85.3 ± 0.40 | 6.56E-12 | 3.02 ± 0.21 | 1.15E-07 | 0.05 ± 0.01 | 2.56E-04 |  |  |  |  |
|  |  | Null | 92.1 ± 0.97 |  | 7.92 ± 0.97 |  | 0 |  | 0 |  |  |  |  |  |
| α1 K219 | P3H3 | WT | 73.1 ± 1.10 | 0.6388 | 26.9 ± 1.10 | 0.6388 | ND | | ND | | | | | |
|  |  | Null | 72.7 ± 1.07 |  | 27.3 ± 1.07 |  | ND |  | ND |  |  |  |  |  |
|  | LH1 | WT | 75.9 ± 1.16 | 2.90E-07 | 24.1 ± 1.16 | 2.90E-07 | ND |  | ND |  |  |  |  |  |
|  |  | Null | 92.7 ± 0.71 |  | 7.28 ± 0.71 |  | ND |  | ND |  |  |  |  |  |
| α2 K219 | P3H3 | WT | 51.6 ± 0.51 | 4.67E-08 | 30.5 ± 0.54 | 0.00259 | 17.1 ± 0.16 | 1.02E-11 | 0.84 ± 0.02 | 5.20E-09 | | | | |
|  |  | Null | 64.1 ± 0.54 |  | 32.5 ± 0.62 |  | 3.27 ± 0.12 |  | 0.14 ± 0.02 |  |  |  |  |  |
|  | LH1 | WT | 48.3 ± 1.30 | 5.04E-08 | 31.4 ± 1.19 | 2.65E-06 | 19.2 ± 0.06 | 2.61E-14 | 1.09 ± 0.07 | 4.93E-08 |  |  |  |  |
|  |  | Null | 87.1 ± 1.94 |  | 12.6 ± 1.86 |  | 0.38 ± 0.08 |  | 0 |  |  |  |  |  |
| α1 K564 | P3H3 | WT | 79.1 ± 1.09 | 0.01211 | 20.1 ± 1.05 | 0.01518 | 0.63 ± 0.04 | 0.00221 | 0.19 ± 0.01 | 1.12E-04 | | | | |
|  |  | Null | 76.7 ± 0.81 |  | 22.3 ± 0.79 |  | 0.77 ± 0.03 |  | 0.27 ± 0.01 |  |  |  |  |  |
|  | LH1 | WT | 80.1 ± 1.61 | 1.05E-06 | 18.8 ± 1.57 | 1.15E-06 | 0.66 ± 0.05 | 4.86E-07 | 0.22 ± 0.02 | 1.83E-04 |  |  |  |  |
|  |  | Null | 96.5 ± 0.19 |  | 3.30 ± 0.19 |  | 0.06 ± 0.01 |  | 0.14 ± 0.01 |  |  |  |  |  |
| α2 K933 | P3H3 | WT | 0.99 ± 0.12 | 2.22E-16 | 93.7 ± 1.15 | 9.40E-12 | 5.34 ± 1.20 | 2.84E-04 | ND | | | | | |
|  |  | Null | 85.6 ± 0.16 |  | 13.6 ± 0.13 |  | 0.81 ± 0.08 |  | ND |  |  |  |  |  |
|  | LH1 | WT | 0.52 ± 0.05 | 0 | 91.6 ± 0.45 | 2.04E-14 | 7.92 ± 0.48 | 5.00E-08 | ND |  |  |  |  |  |
|  |  | Null | 99.2 ± 0.12 |  | 0.75 ± 0.12 |  | 0 |  | ND |  |  |  |  |  |
|  |  |  | Lys-Lys (%) |  | P value |  | Lys-Hyl (%) |  | P value |  | Hyl-Hyl (%) |  | P value |  |
| α1 K918/K930 | P3H3 | WT | 11.3 ± 0.29 | | 1.14E-06 | | 12.7 ± 0.58 | | 0.13635 | | 76.0 ± 0.75 | | 5.17E-10 | |
|  |  | Null | 90.5 ± 8.07 |  |  |  | 7.43 ± 6.15 |  |  |  | 2.05 ± 1.94 |  |  |  |
|  | LH1 | WT | 7.63 ± 0.41 |  | 3.32E-13 |  | 11.3 ± 0.41 |  | 3.26E-09 |  | 81.0 ± 0.75 |  | 3.06E-12 |  |
|  |  | Null | 99.2 ± 0.63 |  |  |  | 0.30 ± 0.08 |  |  |  | 0.51 ± 0.60 |  |  |  |

Values are given as means ± S.D. (n ≥ 3). Biological replicates were n = 4 for all tissues and genotypes.
Lys + Hyl + GHL + GGHL = 100 % and Lys-Lys + Lys-Hyl + Hyl-Hyl = 100 %. Values of modified and unmodified hydroxylysines and unmodified lysine and were obtained using mass spectrometry. Note: Lys; unmodified lysine, Hyl; unmodified hydroxylysine, GHL; galactosyl hydroxylysine, GGHL; glucosylgalactosyl hydroxylysine, Lys-Lys; unmodified lysine and unmodified lysine, Lys-Hyl; unmodified lysine and unmodified hydroxylysine, Hyl-Hyl; unmodified hydroxylysine and unmodified hydroxylysine

**Table 4:** Comparison of lysine post-translational modifications of skin type I collagen at individual sites.

|  |  |  | Lys (%) | P value | Hyl (%) | P value | GHL (%) | P value | GGHL (%) | P value |  |  |  |  |
| --- | --- | --- | --- | --- | --- | --- | --- | --- | --- | --- | --- | --- | --- | --- |
| $\alpha 1$ K87 | P3H3 | WT | 15.7 ± 13.3 | 9.78E-05 | 2.35 ± 0.24 | 1.01E-05 | 7.35 ± 1.08 | 1.03E-04 | 74.6 ± 12.3 | 4.49E-05 | | | | |
| | | Null | 77.8 ± 3.06 | | 10.3 ± 1.15 | | 2.30 ± 0.28 | | 9.65 ± 1.78 | | | | | |
| | LH1 | WT | 8.28 ± 5.34 | 4.34E-08 | 2.06 ± 0.37 | 1.59E-04 | 6.83 ± 0.36 | 2.77E-08 | 82.8 ± 5.18 | 6.40E-08 | | | | |
| | | Null | 98.9 ± 0.12 | | 0.47 ± 0.78 | | 0.20 ± 0.01 | | 0.42 ± 0.03 | | | | | |
| $\alpha 2$ K87 | P3H3 | WT | 6.41 ± 3.13 | 2.95E-08 | 93.6 ± 3.13 | 2.95E-08 | ND | | ND | | | | | |
| | | Null | 73.0 ± 1.92 | | 27.0 ± 1.92 | | ND | | ND | | | | | |
| | LH1 | WT | 2.40 ± 0.28 | 3.33E-15 | 97.6 ± 0.28 | 3.33E-15 | ND | | ND | | | | | |
| | | Null | 98.0 ± 0.24 | | 2.04 ± 0.24 | | ND | | ND | | | | | |
| $\alpha 1$ K99 | P3H3 | WT | 76.5 ± 2.31 | 3.83E-05 | 20.4 ± 1.43 | 1.27E-05 | 2.86 ± 0.91 | 0.00916 | 0.32 ± 0.11 | 0.08426 | | | | |
| | | Null | 91.0 ± 1.40 | | 7.68 ± 1.31 | | 1.11 ± 0.12 | | 0.19 ± 0.06 | | | | | |
| | LH1 | WT | 78.4 ± 0.82 | 1.21E-08 | 17.5 ± 1.06 | 1.46E-07 | 3.54 ± 0.23 | 4.29E-07 | 0.62 ± 0.05 | 9.04E-06 | | | | |
| | | Null | 96.5 ± 0.26 | | 2.70 ± 0.13 | | 0.64 ± 0.11 | | 0.18 ± 0.03 | | | | | |
| $\alpha 1$ K174 | P3H3 | WT | 66.7 ± 4.73 | 3.38E-04 | 32.2 ± 4.34 | 2.82E-04 | 1.16 ± 0.44 | 0.00656 | 0 | 0 | | | | |
| | | Null | 84.2 ± 0.84 | | 15.5 ± 0.79 | | 0.24 ± 0.10 | | 0 | | | | | |
| | LH1 | WT | 63.1 ± 0.50 | 3.00E-11 | 34.4 ± 0.50 | 5.13E-11 | 1.88 ± 0.06 | 9.56E-09 | 0.53 ± 0.02 | 9.22E-09 | | | | |
| | | Null | 92.1 ± 0.07 | | 7.69 ± 0.11 | | 0.24 ± 0.04 | | 0 | | | | | |
| $\alpha 2$ K174 | P3H3 | WT | 40.6 ± 13.1 | 2.16E-04 | 4.86 ± 1.24 | 0.48365 | 50.6 ± 10.6 | 9.59E-05 | 3.94 ± 1.76 | 0.00418 | | | | |
| | | Null | 92.6 ± 1.27 | | 5.40 ± 0.73 | | 2.00 ± 1.21 | | 0 | | | | | |
| | LH1 | WT | 22.7 ± 1.17 | 4.31E-11 | 7.89 ± 0.80 | 3.12E-04 | 60.0 ± 0.79 | 9.10E-12 | 9.47 ± 0.48 | 2.07E-08 | | | | |
| | | Null | 92.9 ± 0.58 | | 4.59 ± 0.39 | | 2.35 ± 0.23 | | 0.20 ± 0.02 | | | | | |
| $\alpha 1$ K219 | P3H3 | WT | 86.2 ± 1.55 | 0.00779 | 13.8 ± 1.55 | 0.00779 | ND | | ND | | | | | |
| | | Null | 90.0 ± 1.11 | | 10.0 ± 1.11 | | ND | | ND | | | | | |
| | LH1 | WT | 85.3 ± 0.48 | 1.67E-07 | 14.7 ± 0.48 | 1.67E-07 | ND | | ND | | | | | |
| | | Null | 95.1 ± 0.54 | | 4.91 ± 0.54 | | ND | | ND | | | | | |
| $\alpha 2$ K219 | P3H3 | WT | 55.1 ± 7.05 | 0.00133 | 34.0 ± 4.12 | 0.0033 | 0.62 ± 0.21 | 0.00161 | 10.3 ± 2.77 | 5.15E-04 | | | | |
| | | Null | 75.2 ± 0.91 | | 23.9 ± 1.09 | | 0.03 ± 0.04 | | 0.88 ± 0.25 | | | | | |
| | LH1 | WT | 50.3 ± 0.79 | 6.51E-10 | 34.7 ± 0.92 | 2.31E-08 | 0.91 ± 0.13 | 1.70E-05 | 13.9 ± 0.63 | 1.42E-08 | | | | |
| | | Null | 85.4 ± 0.65 | | 13.7 ± 0.64 | | 0.08 ± 0.02 | | 0.80 ± 0.09 | | | | | |
| $\alpha 1$ K564 | P3H3 | WT | 77.3 ± 1.20 | 0.7026 | 20.3 ± 1.06 | 0.33603 | 1.72 ± 0.13 | 0.07732 | 0.70 ± 0.12 | 0.13392 | | | | |
| | | Null | 77.6 ± 0.79 | | 19.7 ± 0.65 | | 1.93 ± 0.14 | | 0.85 ± 0.13 | | | | | |
| | LH1 | WT | 78.9 ± 0.82 | 1.20E-07 | 18.4 ± 0.58 | 4.12E-08 | 2.00 ± 0.19 | 7.87E-06 | 0.69 ± 0.10 | 3.30E-04 | | | | |
| | | Null | 92.4 ± 0.47 | | 6.76 ± 0.35 | | 0.55 ± 0.07 | | 0.28 ± 0.05 | | | | | |
| $\alpha 2$ K933 | P3H3 | WT | 0 | 3.58E-11 | 93.1 ± 0.37 | 2.12E-09 | 6.86 ± 2.37 | 0.00117 | ND | | | | | |
| | | Null | 86.7 ± 1.56 | | 13.3 ± 1.56 | | 0 | | ND | | | | | |
| | LH1 | WT | 0 | 5.55E-16 | 93.0 ± 1.23 | 7.36E-12 | 7.02 ± 1.23 | 2.65E-05 | ND | | | | | |
| | | Null | 97.9 ± 0.28 | | 2.05 ± 0.28 | | 0 | | ND | | | | | |
|  |  |  | Lys-Lys (%) |  | P value |  | Lys-Hyl (%) |  | P value |  | Hyl-Hyl (%) |  | P value |  |
| $\alpha 1$ K918/K930 | P3H3 | WT | 1.57 ± 0.73 | | 5.07E-07 | | 8.11 ± 2.66 | | 0.09689 | | 90.3 ± 3.17 | | 3.67E-07 | |
| | | Null | 88.9 ± 7.73 | | | | 5.16 ± 1.41 | | | | 5.99 ± 6.36 | | | |
| | LH1 | WT | 0.29 ± 0.21 | | 1.11E-16 | | 4.00 ± 0.18 | | 4.47E-08 | | 95.7 ± 0.21 | | 3.33E-16 | |
| | | Null | 99.0 ± 0.06 | | | | 0.78 ± 0.07 | | | | 0.22 ± 0.12 | | | |

Values are given as means ± S.D. Biological replicates were n = 4 for all tissues and genotypes.
Lys + Hyl + GHL + GGHL = 100 % and Lys-Lys + Lys-Hyl + Hyl-Hyl = 100 %. Values of modified and unmodified hydroxylysines and unmodified lysine and were obtained using mass spectrometry. Note: Lys; unmodified lysine, Hyl; unmodified hydroxylysine, GHL; galactosyl hydroxylysine, GGHL; glucosylgalactosyl hydroxylysine, Lys-Lys; unmodified lysine and unmodified lysine, Lys-Hyl; unmodified lysine and unmodified hydroxylysine, Hyl-Hyl; unmodified hydroxylysine and unmodified hydroxylysine

### Qualitative and quantitative characterization of skin type V collagen from WT, P3H3 null and LH1 null mice

Type V collagen is heavily lysyl hydroxylated and *O*-glycosylated and abundant in skin compared to tendon and bone [33]. We isolated type V collagen from skin of P3H3 and LH1 null mice and subjected it to gel migration analysis. Although type V collagen from LH1 null mice did not show a clear difference in gel migration however, type V collagen from P3H3 null mice appeared to migrate faster compared to WT (Figure 6A). To confirm these observations, we identified the level of PTMs in both P3H3 null and LH1 null type V collagens by AAA. The ratio in prolyl hydroxylations is slightly different between control animals from the P3H3 and LH1 mouse strains (Figure 6B). One potential explanation is that the analyses were done on skin from 2∼5-month-old and 10-week-old mice for P3H3 and LH1 mice, respectively (more detailed information in the methods section). Similar to type I collagen, neither P3H3 nor LH1 null mice had changes in proline hydroxylations (prolyl 3- and 4-hydroxylation). Surprisingly, P3H3 null mice, but not LH1 null mice, had reduced levels of lysyl hydroxylation in type V collagen isolated from skin (Figure 6B and Table 1) however the occupancy of *O*-glycosylation on hydroxylysine was not changed (Table 2). The reduced lysyl hydroxylation in P3H3 null mice influenced the thermal stability of type V collagen and CD melting curves showed only one of the two thermal transitions seen in WT (Figure 6C). Since lysine residues at the Yaa position of collagenous Gly-Xaa-Yaa triplets are extensively glycosylated in type V collagen [33], site-specific characterization of lysine modifications was difficult due to missed cleavage at hydroxylysine glycosides by trypsin [34]. We were able to analyze two sites, α1(V) K84 and α2(V) K87 (Figure 7 and Table 5), that are involved in crosslink formation [35]. At both sites, the level of GGHL was decreased and the magnitude of reduction of GGHL corresponds to that of the increased unmodified lysines in the absence of P3H3, however this change at α1(V) K84 was not significant statistically (Table 5). This suggests P3H3 could play an important role in lysyl hydroxylation and/or subsequent *O*-glycosylation at the site of crosslink formation consistently. In LH1 null mice, there was a marginal change at α1(V) K84, however, α2(V) K87 was clearly affected. Potential explanation is that the α2-chain of type V collagen is classified as an A-clade chain, which includes both the α1- and α2-chain of type I collagen, whereas the α1-chain of type V collagen belongs to B-clade [36, 37]. LH1 seems to hydroxylate the α2(V) K87 preferentially. Nevertheless, the ratio of two α1-chains and one α2-chain in type V collagen could hide the effect caused by impaired LH1 activity and not show any distinct difference in type V collagen between WT and LH1 null observed in Figure 6B. In summary, P3H3 is required for proper lysyl hydroxylation of type I and type V collagen whereas LH1 is dispensable for lysyl hydroxylation in skin type V collagen.

**Figure 6.**
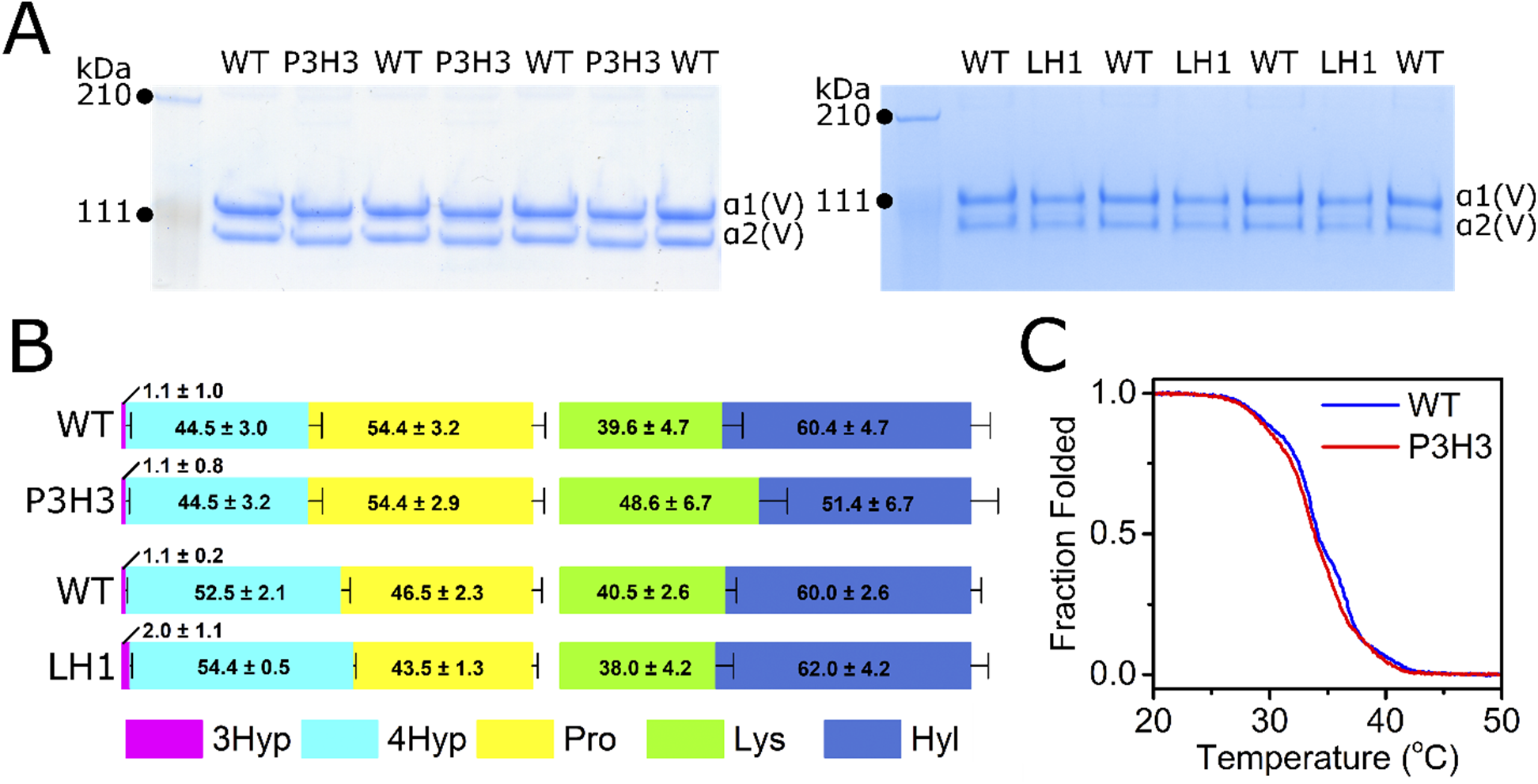
Characterization of skin type V collagen from P3H3 and LH1 null mice. (A) SDS-PAGE analysis of purified pepsin treated skin type V collagen of WTs, P3H3 and LH1 null mice. The final purified material in the presence of reducing agent was run running on a NuPAGE 3 – 8 % Tris-Acetate gel stained with GelCode Blue Stain Reagent. For both SDS-PAGE analysis of P3H3 and LH1, each genotype had three biological replicates since each lane in gel was loaded by independently prepared collagen from tissue. (B) The ratio of post-translational modifications in proline (3Hyp + 4Hyp + Pro = 100) and lysine (Lys + Hyl = 100) in skin type V collagen of WTs, P3H3 null and LH1 null are demonstrated as bar graphs which are generated by values from Table 1. Values of amino acids were obtained using amino acid analysis and biological replicates were n = 17 for P3H3, n = 3 for LH1 WT and n = 6 for LH1 null. The numbers in the graphs indicate the mean ± S.D. of individual amino acids and P values obtained by statistical analyses are in Table 1 [3Hyp(magenta); 3-hydroxyproline, 4Hyp (cyan); 4-hydroxyproline n, Pro (yellow); unmodified proline, Lys (green); unmodified lysine, Hyl (blue); hydroxylysine]. (C) Thermal stability of skin type V collagen from P3H3 WT (blue) and null (red) was monitored by CD at 221 nm in 0.05 M acetic acid, and the rate of heating was 10 °C/h. Biological replicates of each curve were n = 2 for both WT and P3H3 null. In figures, P3H3 and LH1 indicate P3H3 null and LH1 null, respectively. Tissues were collected from 2∼5-month-old mice.

**Figure 7.**
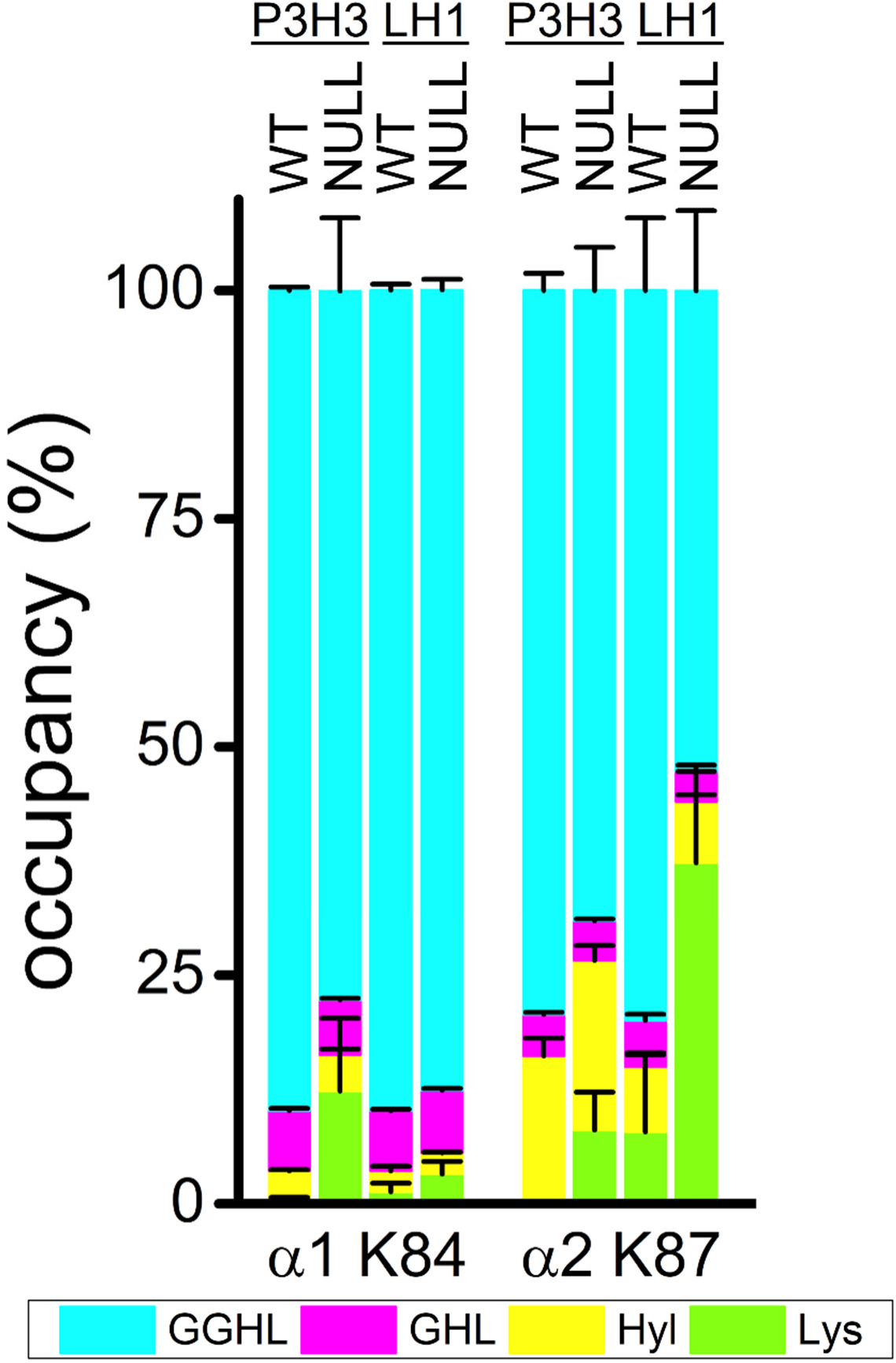
Summary of lysine post-translational modifications of skin type V collagen in individual sites. Bar graphs represent the occupancy of lysine modifications in individual lysyl hydroxylation sites of skin type V collagen of WTs, P3H3 null and LH1. Individual values [GGHL (cyan); glucosylgalactosyl hydroxylysine, GHL (magenta); galactosyl hydroxylysine, Hyl (yellow); unmodified hydroxylysine, Lys (green); unmodified lysine] correspond to Table 5. Values of modified and unmodified hydroxylysines and unmodified lysine were obtained using mass spectrometry and biological replicates were n =3 for P3H3 WT and null, n ≥ 3 for LH1 WT and n = 4 for LH1 null. α1, α2 and K + numbers indicate that α1 and α2 chain of type V collagen and residue number from the first residue of triple helical domain.

**Table 5:** Comparison of lysine post-translational modifications of skin type V collagen in individual sites.

|  |  |  | Lys (%) | P value | Hyl (%) | P value | GHL (%) | P value | GGHL (%) | P value |
| --- | --- | --- | --- | --- | --- | --- | --- | --- | --- | --- |
| $\alpha 1$ K84 | P3H3 | WT | $0.63 \pm 0.04$ | 0.06323 | $2.97 \pm 0.09$ | 0.05873 | $6.61 \pm 0.37$ | 0.23762 | $89.8 \pm 0.36$ | 0.05822 |
| | | Null | $12.3 \pm 7.94$ | | $3.95 \pm 0.65$ | | $6.01 \pm 0.56$ | | $77.6 \pm 8.01$ | |
| | LH1 | WT | $1.33 \pm 0.92$ | 0.05917 | $2.28 \pm 0.48$ | 0.92243 | $6.56 \pm 0.17$ | 0.06623 | $89.9 \pm 0.66$ | 0.01655 |
| | | Null | $3.25 \pm 1.38$ | | $2.30 \pm 0.12$ | | $6.88 \pm 0.21$ | | $87.6 \pm 1.18$ | |
| $\alpha 2$ K87 | P3H3 | WT | 0 | 0.02794 | $16.2 \pm 1.95$ | 0.17434 | $4.55 \pm 0.22$ | 0.26084 | $79.3 \pm 1.88$ | 0.0248 |
| | | Null | $8.07 \pm 4.14$ | | $18.6 \pm 1.59$ | | $4.55 \pm 0.17$ | | $69.0 \pm 4.71$ | |
| | LH1 | WT | $7.87 \pm 8.46$ | 0.00938 | $7.13 \pm 1.51$ | 0.61402 | $5.07 \pm 0.65$ | 0.01839 | $79.9 \pm 7.92$ | 0.00827 |
| | | Null | $37.3 \pm 1.00$ | | $6.68 \pm 0.74$ | | $3.33 \pm 0.67$ | | $52.7 \pm 8.79$ | |

Values are given as means ± S.D. Biological replicates were n =3 for P3H3 WT and null, n ≥ 3 for LH1 WT and n = 4 for LH1 null.
Lys + Hyl + GHL + GGHL = 100 %. Values of modified and unmodified hydroxylysines and unmodified lysine were obtained using mass spectrometry.
Note: Lys; unmodified lysine, Hyl; unmodified hydroxylysine, GHL; galactosyl hydroxylysine, GGHL; glucosylgalactosyl hydroxylysine

### Skin analysis in P3H3 null mice

Consistent with skin type V collagen analysis, and as reported previously [18], we found defects in skin from P3H3 null mice. Masson’s Trichrome staining shows less collagen content and the thickness and ratio were altered in the dermis and hypodermis (Figure 8A – C). Electron microscopy showed that the average diameter of collagen fibrils is similar between P3H3 null (84.9 ± 35.6 nm) and WT (85.1 ± 25.9 nm) (Figure 8D), however the distribution of fibril diameters was broader in P3H3 null (Figure 8E) and this was also reported in LH1 null skin [26]. In summary, a precise number of PTMs in the rER is required to maintain an appropriate ultrastructure in collagen rich tissues.

**Figure 8.**
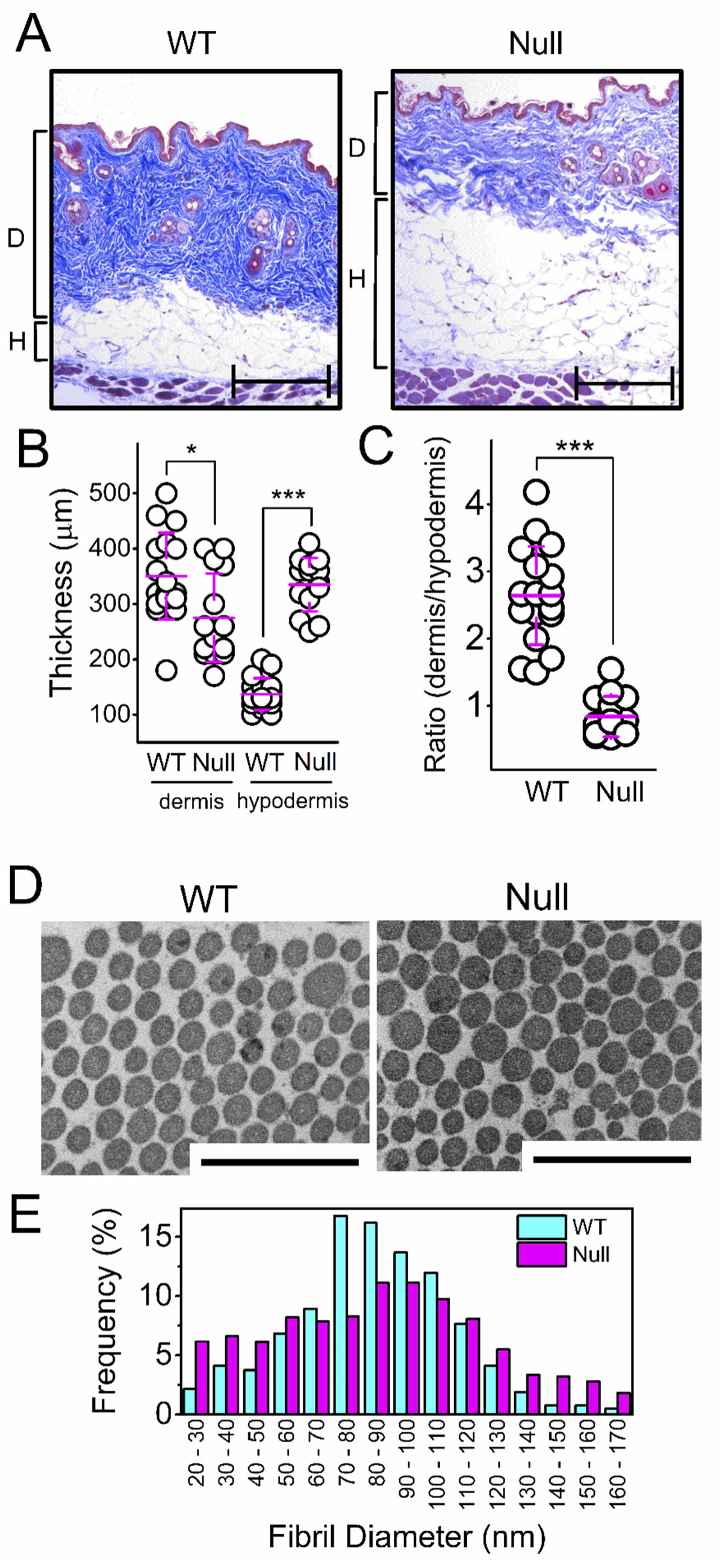
Skin analysis of P3H3 null mice. (A) Masson’s Trichrome stain was performed using skin section from 8-month-old P3H3 WT and null mice. D and H indicate dermis and hypodermis corresponding to the blue rich and white rich area, respectively. Scale bars, 200 µm. (B) The thickness of dermis and hypodermis were measured from the images of Masson’s Trichrome stains. The averaged thickness of dermis and hypodermis are 350.6 ± 78.2 µm and 137.1 ± 28.7 µm for WT and 275.0 ± 80.1 µm and 335.7 ± 48.6 µm for P3H3 null. * and *** indicate P < 0.05 (P = 0.013) and P < 0.0005 (P = 1.4988E^-14^), respectively. (C) The ratio between dermis and hypodermis was calculated by the set of thickness including both dermis and hypodermis from Masson’s Trichrome stains. The averaged ratio between dermis and hypodermis are 2.64 ± 0.73 for WT and 0.84 ± 0.30 for P3H3 null. *** indicate P < 0.0005 as P = 1.96408E^-9^. (D) Electron microscopy images display the skin collagen fibrils in 3-month-old P3H3 WT and null mice. Scale bar, 500 nm. (E) The histogram represents the fibril diameter distribution in P3H3 WT (cyan) and null (magenta). The averaged fibril diameter of P3H3 null and WT are 84.9 ± 35.6 nm and 85.1 ± 25.9 nm, respectively. The thickness of collagen fibrils was counted using electron microscopy images.

## Discussions

In the rER, many enzymes and post-translational modifiers interact with molecular chaperones via either strong or weak affinity interaction to improve their functions [38, 39]. In particular, the molecular ensemble for collagen biosynthesis consists of variety of protein–protein interactions [13, 40]. When an interaction is impaired, as found in genetic disorders, the magnitude of the impact depends on what type of protein–protein interaction is disrupted. Prolyl 3-hydroxylase 1 (P3H1), cartilage-associated protein (CRTAP) and cyclophilin B (CypB) form a complex with very tight molecular interactions [41, 42]. In this complex each protein stabilizes the others and CRTAP requires the other two proteins to maintain its solubility [42–44]. As a result, the lack of one molecule in this complex leads to very similar phenotypic abnormalities in osteogenesis imperfecta (OI) [20, 45]. On the other hand, CypB is also identified as a binding partner for LH1. CypB is an ER-resident peptidyl-prolyl cis-trans isomerase and was the first molecule shown to be associated with LH1 [46]. The absence of CypB or disruption of the interaction with LH1 results in the reduction of lysyl hydroxylation in tendon and skin [31, 32], while the amount of lysyl hydroxylation is increased in bone [47]. Additionally, there is a site-specific effect to the level of lysyl hydroxylation in tendon, skin and bone. Therefore CypB was proposed to control lysyl hydroxylation [32]. SC65, a homolog of CRTAP, was suggested to be interacting with P3H3 with relatively strong affinity as indicated by gel filtration chromatography combined with Western blots [48]. In the SC65 null mouse, a reduction of lysyl hydroxylation at α1(I) and α2(I) K87 was found in tissues, whereas these changes did not always correspond to the changes found in P3H3 null mice [18]. Overall, mouse models of LH1 and LH1-associated proteins showed aberrant crosslink formations caused by lysine under hydroxylation at the α1 K87 and α1 K930, therefore three molecules (CypB, SC65 and P3H3) have been proposed to form a complex with LH1 in the rER [48]. However, it is not clear how these molecules interact with each other and what effect one has on other molecules.

To evaluate the correlation between LH1 and one of the LH1-associated proteins, P3H3, we conducted quantitative analyses and directly compared the level of PTMs between WT, P3H3 null and LH1 null mouse tissues. Our results suggest that if P3H3 acts as a LH1 chaperone, this chaperone function is not required for all LH1 sites, as very specific sites related to crosslink formation in type I collagen were affected (Figure 5). Figure 9 represents the magnitudes of change of unmodified lysine residues in individual lysyl hydroxylation sites between three different null mouse models compared to WT. These observations imply that both P3H3 and CypB play important roles for the function of LH1 and that a lack of even one of the components attenuates the amounts of hydroxylysine in the crosslink formation sites. Conversely, other lysyl hydroxylation sites demonstrate very diverse effects between P3H3 null, LH1 null and CypB null mouse tissues. There are specific patterns that are changed in each null mouse model. Modified lysine residues were hardly found in LH1 null, whereas P3H3 null showed normal or slightly increased unmodified lysine residues. In contrast, CypB null showed normal or decreased unmodified lysine residues despite increasing unmodified lysine residues at crosslink formation sites as well as P3H3 and LH1 nulls. Moreover, additional sugar attachments are found at other lysyl hydroxylation sites (e.g. K174 and K219) in the P3H1 null mice model [30] which displays an OI phenotype [49] and CypB null is also a model of recessive type IX OI [47]. This suggests that these proteins do not form a tight molecular complex like the P3H1/CRTAP/CypB complex but more likely a combination of distinct protein–protein interactions. Indeed, this variety of interactions was shown by Western blotting results between LH1 and three LH1 associated protein. The protein level of P3H3 was not abolished but decreased in the absence of CypB and SC65 [32, 48]. Similarly, SC65 protein was affected in CypB null mice [32]. However, the protein level of LH1 was decreased as well as P3H3 in the absence of SC65 whereas the lack of CypB interestingly induced more LH1 protein than WT [32, 48]. Additionally, size exclusion chromatography demonstrated that P3H3 and SC65 were possibly associated in a tight interaction like the P3H1/CRTAP/CypB complex, however neither LH1 nor CypB was a part in this tight interaction [48]. We imagine a very precise molecular interplay is required particularly around crosslink formation sites and this is not simply determined as chaperone effects and/or a complex formation. Collectively, we would like to term this precise mechanism as a “local molecular ensemble”. We looked for the specific binding or enhancer sequences of type I collagen near lysyl hydroxylation sites based on our results (Figure 10). As a previous report suggested [32], the KGH sequence occurs at or near crosslink formation sites to provide preferential interaction sites for the CypB-involved SC65/P3H3 ER complex to facilitate LH1 activity. This hypothesis is possible, but cannot explain the reduction of α2(I) K87, K174 and K933 since the KGH sequence only exist near α2(I) K87 in mouse and does not exist near α2(I) K174. (Figure 10). Interestingly, some ECM proteins (integrins, decorin and SPARC) are likely associated at around K174 of type I collagen [50]. Thus, α2(I) K174 could have important roles since this site is highly sensitive in the absence of P3H3, LH1 and CypB (Figure 9). In type V collagen, both α1(V) K84 and α2(V) K87 are involved in crosslink formation [35] and these site are also the KGH sequence (Figure 10). However, the effects at these sites are not consistent between P3H3 and LH1 null (Figure 7). Further studies are required to elucidate how these four molecules properly distinguish and gather to form a local molecular ensemble at individual lysyl hydroxylation sites at a molecular level. Here we note that a previous report showed the difference in PTMs at α1(V) K87 [18], however the actual residue 87 is arginine instead of lysine as we showed above and this is also confirmed by database (UniProt entry numbers: P20908 for human and O88207 for mouse, NCBI accession number: bovine for XP_024855494).

**Figure 9.**
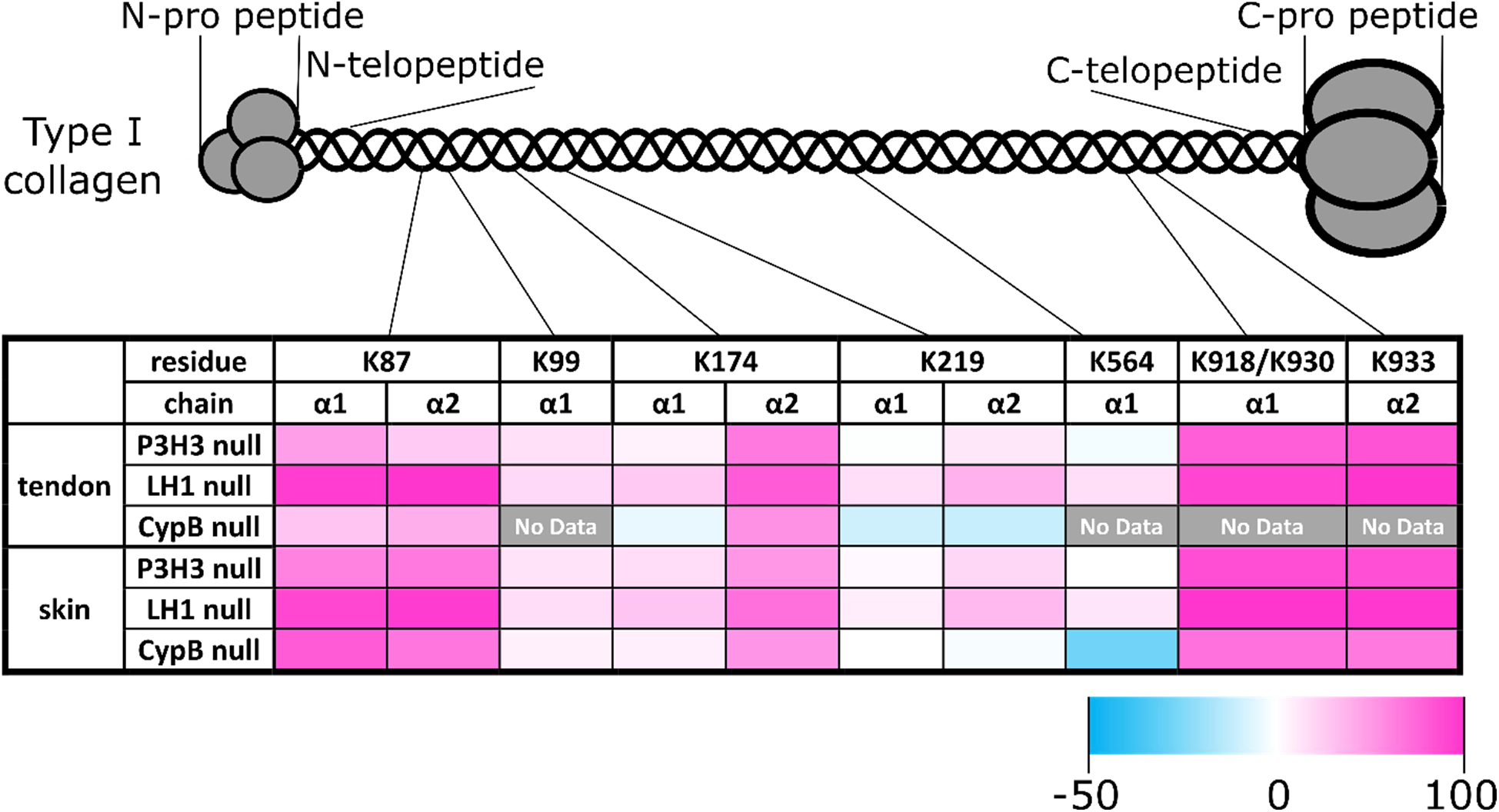
The magnitudes of change of unmodified lysine residues in individual lysyl hydroxylation sites of type I collagen between null mice models compared to WT. Schematic diagram and heat map show the magnitudes of change of unmodified lysine residues in individual lysyl hydroxylation sites. This change was calculated by [the value of unmodified lysine (%) in null mice] – [the value of unmodified lysine (%) in WT mice]. The values in Tables 3 and 4 were used for P3H3 null and LH1 null mice to calculate the changes. The values of unmodified lysine were obtained from reference [31] and [32] for CypB null mice in tendon and skin, respectively.

**Figure 10.**
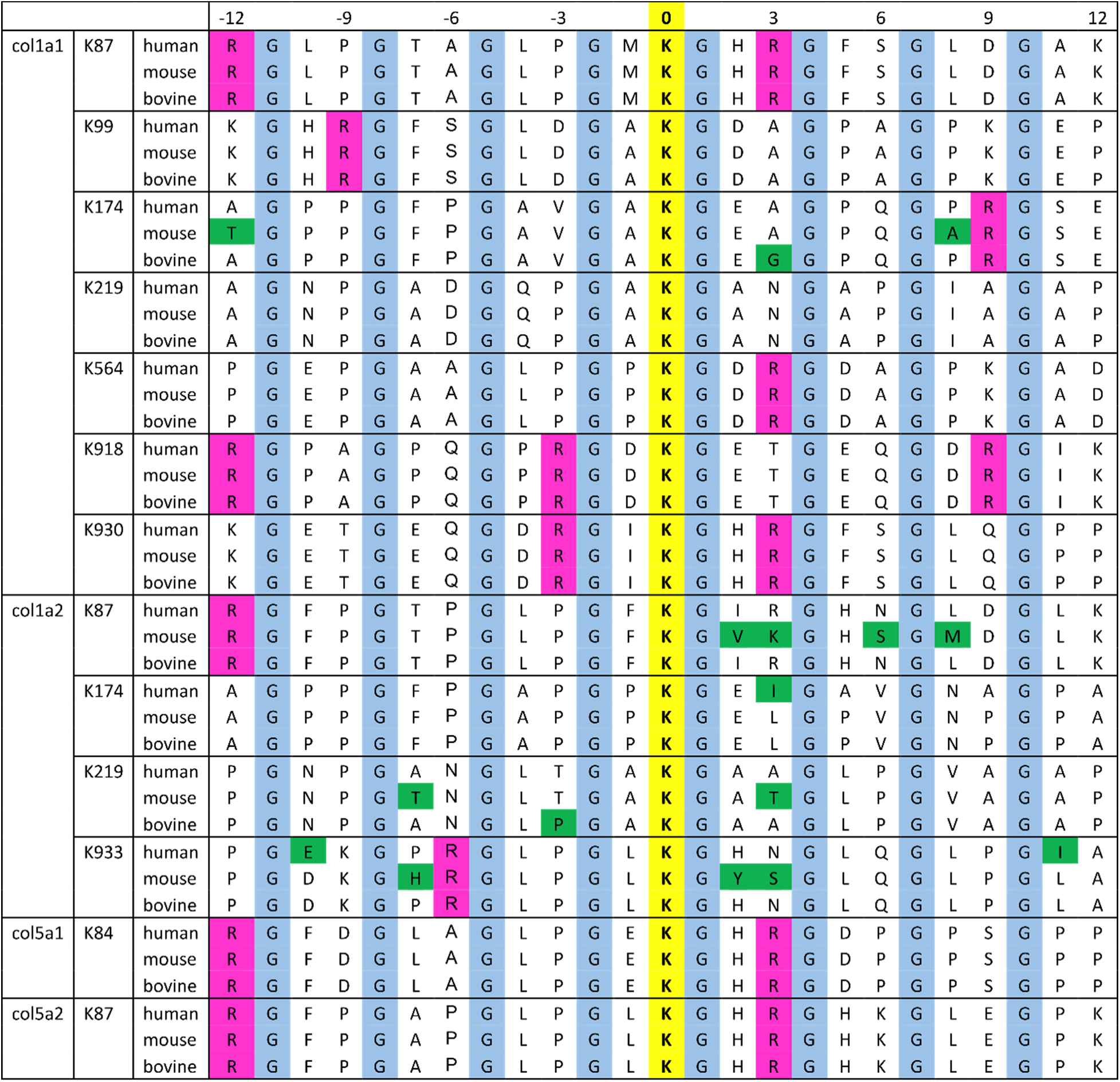
The sequence alignment surrounding lysyl hydroxylation sites in type I and type V collagen between human, mouse and bovine. The sequences are aligned ± 12 residues from the lysine residue which is modified to hydroxylysine and highlighted by yellow with Bold. Glycine residues in GXY repeats and the residues not conserved between human, mouse and bovine are highlighted by cyan and green. Arginine residues in RGXY sequences are also highlighted by magenta because these residues are critical for Hsp47 to bind to collagen triple helices [53]. Uniprot entry numbers are as follows: human COL1A1 (P02452), mouse COL1A1 (P11087), bovine COL1A1 (P02453), human COL1A2 (P08123), mouse COL1A2 (Q01149), bovine COL1A2 (P02465), human COL5A1 (P20908), mouse COL5A1 (O88207), human COL5A2 (P05997) and mouse COL5A2 (Q3U962). NCBI accession numbers are follows: bovine COL5A1 (XP_024855494) and bovine COL5A2 (XP_024835542)

The previous studies of LH1 null and EDS type VIA patients suggested tissue specific and collagen type specific lysyl hydroxylation could exist since different magnitudes of reduction in lysyl hydroxylation between tissues was found [22–25]. Potential explanations were suggested such as the complexity of different collagen types between tissues, a distribution in the expression and/or protein levels of LH isoenzymes or a compensation by two other LH isoenzymes, which could hydroxylate the peptides containing the sequences of hydroxylation sites in triple helices of type I and type IV collagen [9, 16, 51]. Here, we show that the purified type I collagen without telopeptide regions of tendon, skin and bone from LH1 null are differentially modified between tissues (Figure 3). We also find a decrease of total amount of hydroxylysine in type I collagen from P3H3 null mouse tissues, however the decrease was less than in LH1 null mouse tissues (Figure 3). In contrast to LH1 null and P3H3 null, CypB null mice showed the tissue dependent alteration in total amount of hydroxylysine. Hydroxylysine was reduced in type I collagen from tendon and skin while an increase was observed in bone [31, 32, 47]. Therefore, lysyl hydroxylation in triple helical domain of type I collagen is likely reactive by disruption of LH1 and LH1 associated proteins. Contrary to the results found in type I collagen, there was interestingly no significant difference in type V collagen from LH1 null skin [26]. Surprisingly, P3H3 null showed an obvious reduction of lysyl hydroxylation in type V collagen from skin as well as type I collagen (Figures 3 and 6). Considering the results from the lung and kidney of LH3 mutant mice that demonstrate that the amount of hydroxylysine was not changed in type I collagen rich fractions, but was reduced by 30% in type IV and type V collagen rich fractions [24], we suggest that LH1 and LH3 have a substrate specificity for at least type I collagen and type V collagen, respectively. In addition, given that the LH3 mutant mice only showed a decrease of 30% in type IV and type V collagen rich fractions, that type V collagen from P3H3 null showed a decrease of hydroxylysine content and also that the hydroxylysine content of bone type I collagen was decreased in P3H3 null despite increasing in CypB null, we speculate that P3H3 might have a lysyl hydroxylation activity, namely, LH4 like enzyme in the rER. Although P3H3 belongs to the prolyl 3-hydroxylase family, no substrate or enzyme activity have been identified. P3H3 could interact with both LH1 and LH3, however this is unlikely because the phenotype of P3H3 null mice is very mild. Thus, direct prolyl and lysyl hydroxylase activity assays are needed to determine if P3H3 acts as prolyl and/or lysyl hydroxylase. However, attempts to produce necessary quantities of recombinant protein have not succeeded.

In conclusion, P3H3 and LH1 play critical roles to hydroxylate lysine residues in crosslink formation sites in type I collagen whereas they likely have distinct mechanisms to modify other sites in type I collagen and to recognize different collagen types in the rER. Indeed, it is still unclear what the most important factors are to obtain the precise PTMs in collagens: is it the formation of specific protein– protein interactions or the level of expression that leads to the existence of active complexes? Our findings offer new directions for the understanding of lysyl hydroxylation in different tissues and in different collagens and provide a new interpretation of previous findings in LH1 null mice, LH3 null mice and EDS VI patients.

## Experimental procedures

### P3H3 null mice

P3H3 null mice were purchased from Ozgene (Bentley, Australia). Directed knockouts were created in which exon 1 of the mouse *Leprel2* (also called *P3H3*) gene (UniProt entry number: Q8CG70), coding P3H3, were deleted. The procedure is briefly described as follow. The PGK-neo selection cassette flanked by FRT sites was generated by PCR from C57BL/6 genomic DNA and inserted into the downstream of the exon 1 which was flanked by loxP sites. The targeted locus was eliminated using an FLP recombinase and a Cre recombinase. ES cell clones were confirmed by southern hybridization. P3H3 inactivation was verified in mice by DNA preparation from tissues and PCR with primer sets (Fwd: 5’-CTTACCCACACTAGACCCATGTGTC-3’ and Rev: 5’-GTTGCATTCATTAGCCTAGACCCGCTA-3’). PCR was performed at least 30 times and the PCR products for wild type and null were located at around 1,000 and 200 pb, respectively. Tissues were harvested from 2∼5-month-old mice for collagen analysis, 8-month-old mice for skin staining analysis and 3-month-old mice for fibril diameter analysis.

### LH1 null mice

Plod1 knockout mice were produced as described earlier [26] using in-frame insertion of lacZ-neo cassette into exon 2. Mice were further backcrossed into C57BL/6N background. Genotyping was done by PCR [26] and tissues were harvested from 10-week-old mice for collagen analysis.

### Western Blotting

A kidney was extracted from wild type, heterozygous and null P3H3 mouse and homogenized using T-PER (Thermo Fisher Scientific) containing protease inhibitors at 4 °C. After centrifugation, soluble proteins in the extract were mixed with NuPAGE LDS sample buffer with reducing agents. These protein solutions were separated by Bolt 4-12% Bis-Tris Plus (Life Technology) and electrotransferred onto PVDF membranes. Antibodies were incubated to detect the specific protein after the membranes were blocked in PBS solution containing 5% (w/v) skim milk. All proteins were detected by alkaline phosphatase developed with 5-bromo-4-chloro-3-indolyl phosphate and Nitro blue tetrazolium. Rabbit polyclonal antibody against P3H3 (*LEPREL2*) and rabbit polyclonal antibody against GAPDH were purchased from Proteintech (16023-1-AP) and Sigma-Aldrich (PLA0125), respectively. Alkaline phosphatase-conjugated anti-rabbit IgG (A9919: Sigma-Aldrich) was used as secondary antibody. Western blotting was performed three times using independently prepared three independent kidneys.

### X-ray scan

X-rays were performed at least three times on independently prepared adult mice using a Faxitron cabinet instrument (model #43855B) made by Hewlett Packard. Voltages and exposure times were optimized for best resolution images.

### Collagen Extraction from Tissues

Tendon, skin and bone were taken from adult mice. All procedures were performed at 4 °C. Tendon and skin were incubated in excess volume of 0.1 M acetic acid with shaking for several hours. Ground bone in liquid nitrogen was incubated in 1.0 M acetic acid containing 0.05 M EDTA. Pepsin was added to a final concentration of 0.25 mg/mL and tissues were digested overnight. For bone, pepsin digestion and decalcification were taken three days. The solutions were centrifuged to remove insoluble material, and then NaCl was added to a final concentration of 0.7 M to precipitate collagens and the solution was incubated overnight. Precipitates were collected by centrifugation at 13,000 rpm for 15 min and resuspended in 0.1 M acetic acid. This solution was enriched type I collagen from tendon and bone and was dialyzed against 0.1 M acetic acid to remove remaining NaCl. For skin, the solution was dialyzed in excess volume of 0.1 M Tris/HCl containing 1.0 M NaCl, pH 7.8 and then NaCl was added to a final concentration of 1.8 M to remove type III collagen. This solution was centrifuged at 13,000 rpm for 30 min and additional NaCl was added to a final concentration of 2.4 M to the supernatant. After incubating overnight, the solution was centrifuged at 13,000 rpm for 30 min. The pellets containing skin type I collagen was resuspended in 0.1 M acetic acid and dialyzed against 0.1 M acetic acid to remove remaining NaCl. Skin type V collagen was extracted using the supernatant of 0.7 M NaCl precipitation and additional NaCl was added to a final concentration of 4.0 M. After incubating overnight, the solution was ultra-centrifuged at 30,000 rpm for 30 min. The pellets containing skin type I collagen was resuspended in 0.1 M acetic acid and dialyzed against 0.1 M acetic acid to remove remaining NaCl.

### Circular Dichroism

Circular dichroism spectra were recorded on an AVIV 202 spectropolarimeter (AVIV Biomedical, Inc., Lakewood, NJ) using a Peltier thermostatted cell holder and a 1-mm path length rectangular quartz cell (Starna Cells Inc., Atascadero, CA). The temperature scanning curves were monitored at 221 nm with 10 °C/h scan rate. All curves were the average of at least three and two independent measurements using independently prepared collagen from tissue for type I and type V collagen, respectively.

### Amino Acid Analysis

Acid hydrolysis was performed in 6 x 50-mm Pyrex culture tubes placed in Pico Tag reaction vessels fitted with a sealable cap (Eldex Laboratories, Inc., Napa, CA). Samples were placed in culture tubes, dried in a SpeedVac (GMI, Inc. Albertville, MN), and then placed into a reaction vessel that contained 250 ml of 6 M HCl (Pierce) containing 2% phenol (Sigma-Aldrich). The vessel was then purged with argon gas and evacuated using the automated evacuation workstation Eldex hydrolysis/derivatization workstation (Eldex Laboratories, Inc.). Closing the valve on the Pico Tag cap maintained the vacuum during hydrolysis at 105 °C for 24 h. The hydrolyzed samples were then dried in a Savant SpeedVac. The dried samples were dissolved in 100 ml of 0.02 M HCl containing an internal standard (100 µM norvaline; Sigma). Analysis was performed by ion exchange chromatography with postcolumn ninhydrin derivatization and visible detection (440 nm/570 nm) with a Hitachi L-8800A amino acid analyzer (Hitachi High Technologies America, Inc., San Jose, CA) running the EZChrom Elite software (Scientific Software, Inc., Pleasanton, CA). Three technical replicates were performed in each analysis.

### Glycosylation analysis

Glycosylation of hydroxylysine was estimated by LC–MS after alkaline hydrolysis as described previously [52]. In brief, type I and type V collagen samples were subjected to alkaline hydrolysis (2 N NaOH, 110 °C for 20 h under N2) after adding stable isotope-labeled collagen as an internal standard. The alkaline hydrolysates were neutralized with 30% acetic acid and then desalted using a mixed-mode cation-exchange sorbent (Oasis MCX; Waters, Milford, MA). Unmodified and glycosylated hydroxylysines were quantitated by LC–MS in multiple reaction monitoring mode using a 3200 QTRAP hybrid triple quadrupole/linear ion trap mass spectrometer (AB Sciex, Foster City, CA) coupled to an Agilent 1200 Series HPLC system (Agilent Technologies, Palo Alto, CA) with a ZIC-HILIC column (3.5 µm particle size, L × I.D. 150 mm × 2.1 mm; Merck Millipore, Billerica, MA).

### Site-Specific Characterization of Lysine PTMs

Site occupancy of lysine PTMs was estimated by LC– MS after protease digestion as described previously [32]. In brief, type I collagen samples were digested with trypsin (Promega, Madison, WI) or with collagenase from *Grimontia hollisae* (Wako Chemicals, Osaka, Japan) and pepsin (Sigma-Aldrich) after heat denaturation at 60 °C for 30 min. On the other hand, type V collagen samples were separated by SDS-PAGE, and the region containing α1(V) and α2(V) were subjected to in-gel digestion with trypsin. The protease digests of type I and type V collagen were analyzed by LC–MS on a maXis II quadrupole time-of-flight mass spectrometer (Bruker Daltonics, Bremen, Germany) coupled to a Shimadzu Prominence UFLC-XR system (Shimadzu, Kyoto, Japan). Site occupancy of each modification site was calculated using the peak area ratio of monoisotopic extracted ion chromatograms of peptides containing the respective molecular species. α1(I) K918/K930 and other sites were analyzed using the collagenase/pepsin digests and the trypsin digests, respectively.

### Masson’s Trichrome stain

OHSU histology core facility performed sectioning and staining skin sample. Ventral skin tissues (n = 3) were fixed in 4% paraformaldehyde for 24 hours at 4°C. The fixed tissues were dehydrated using an ethanol gradient, cleared in xylene, and embedded in paraffin wax, after which 5 µm sections were cut using a microtome. The tissue sections were stained with Masson’s trichrome for collagen fiber analysis. Regions of interest (n = 2 - 4) were selected per image (n = 5) and thickness of dermis and hypodermis were measured. Length was normalized by each scale bar.

### Electron Microscopy Analysis of skin

The three independently prepared skin from P3H3 null and WT mice was fixed in 1.5 % glutaraldehyde/1.5 % paraformaldehyde (Electron Microscopy Sciences) in Dulbecco’s serum-free media (SFM) containing 0.05 % tannic acid. The samples were rinsed in SFM, post-fixed in 1 % OsO4 then dehydrated in a graded series of ethanol to 100 %, rinsed in propylene oxide and infiltrated in Spurr’s epoxy. Samples were polymerized at 70 °C for 18 hours. Ultra-thin sections (∼80 nm) were cut on a Leica EM UC7 ultramicrotome and mounted on formvar-coated, copper palladium 1×2 mm slot grids. Sections were stained in saturated uranyl acetate followed by lead citrate and photographed using an AMT 2K − 2K side entry camera (AMT, Woburn, MA) mounted on a FEI G2 transmission electron microscope operated at 120 kV and at least four images were taken per sample.

### Fibril diameter measurement

The number and cross-sectional area of fibrils were measured using the Fiji software (ImageJ). Regions of interest were selected from the images of WT (n = 13) and P3H3 null (n = 17) and fibril areas were measured by counting the pixels per fibril, which accounts for μm^2^. Spatial calibration for diameter measurements was applied against each scale bar. The numbers were counted as 1845 and 3616 fibrils for P3H3 WT and null.

### Statistical analyses

For comparisons between two groups, we performed one-way ANOVA to determine whether differences between groups are significant using ORIGIN Pro ver. 9.1 (OriginLab Corp., Northampton, MA). The P value less than 0.05 was considered statistically significant.

## Acknowledgements

We thank to the Analytical Core Facility of Shriners Hospitals for Children in Portland for amino acid analysis and also to the OHSU histology core facility for sectioning and staining skin sample. We also thank Douglas B. Gould (Department of Ophthalmology, University of California, San Francisco) for critical reading of the manuscript and providing valuable comments. This study was supported by grants from Shriners Hospital for Children (85100 and 85500 to HPB) and by the Academy of Finland Project Grant 296498 and Center of Excellence 2012-2017 Grant 251314 (JM), the S. Jusélius Foundation (JM), and the Jane and Aatos Erkko Foundation (JM).

## Author contributions

YI and HPB were responsible for the overall design of the study. YI, YT, KZ, NM, AS, OS, ST and DRK conducted and analyzed experiments. YI, YT, KZ, NM, AS, OS, ST, DRK, PH, KM, JM and HPB provided essential material, reviewed and discussed the results. YI and HPB wrote the main manuscript text. All authors were involved in editing the manuscript.

## Additional information

Competing Interests: The authors declare that they have no competing interests related to this work.

